# A flagellate-to-amoeboid switch in the closest living relatives of animals

**DOI:** 10.1101/2020.06.26.171736

**Authors:** Thibaut Brunet, Marvin Albert, William Roman, Danielle C. Spitzer, Nicole King

## Abstract

The evolution of different cell types was a key process of early animal evolution^1–3^. Two fundamental cell types, epithelial cells and amoeboid cells, are broadly distributed across the animal tree of life^4,5^ but their origin and early evolution are unclear. Epithelial cells are polarized, have a fixed shape and often bear an apical cilium and microvilli. These features are shared with choanoflagellates – the closest living relatives of animals – and are thought to have been inherited from their last common ancestor with animals^1,6,7^. The deformable amoeboid cells of animals, on the other hand, seem strikingly different from choanoflagellates and instead evoke more distantly related eukaryotes, such as diverse amoebae – but it has been unclear whether that similarity reflects common ancestry or convergence^8^. Here, we show that choanoflagellates subjected to spatial confinement differentiate into an amoeboid phenotype by retracting their flagella and microvilli, generating blebs, and activating myosin-based motility. Choanoflagellate cell crawling is polarized by geometrical features of the substrate and allows escape from confined microenvironments. The confinement-induced amoeboid switch is conserved across diverse choanoflagellate species and greatly expands the known phenotypic repertoire of choanoflagellates. The broad phylogenetic distribution of the amoeboid cell phenotype across animals^9–14^ and choanoflagellates, as well as the conserved role of myosin, suggests that myosin-mediated amoeboid motility was present in the life history of their last common ancestor. Thus, the duality between animal epithelial and crawling cells might have evolved from a temporal phenotypic switch between flagellate and amoeboid forms in their single-celled ancestors^3,15,16^.

---

Amoeboid cell motility is central to several key aspects of animal biology, including development^17,18^, immunity^13,14^, and wound healing^19^. Nonetheless, the origin of animal amoeboid cells has remained mysterious^8^, in part because the closest living relatives of animals, the choanoflagellates^20,21^, have been thought to exist solely in a flagellate form (except while encysted^22^). The nature of the protozoan ancestor of animals was a matter of debate as early as the 19^th^ century, when the relationship between animals and choanoflagellates was still unknown. Haeckel originally proposed in 1874 that animals descended from amoebae that evolved coloniality and later acquired cilia^23^. In contrast, Metschnikoff proposed in 1886 that animals originated from colonies of flagellated cells resembling modern choanoflagellates^24^, a view that was incorporated into later interpretations of Haeckel’s Gastraea hypothesis^25^. An intermediate view^26^ was inspired by the discovery of protozoans such as *Naegleria*, which can alternate between a flagellate and an amoeboid form^27^. Under this scenario, the protozoan ancestor of animals may have already contained the genetic programs required for the evolution of differentiated crawling cells (such as sponge archeocytes, cnidarian amoebocytes, and vertebrate white blood cells) and flagellated cells (such as sperm cells and epithelial cells, which often retain an apical cilium/flagellum and/or microvilli^26^).

However, subsequent phylogenetic analyses revealed that protozoans known to alternate between flagellate and amoeboid forms (such as *Naegleria*) belong to branches of the tree of life that are far removed from animals^28^, making their relevance to animal origins uncertain. A single report of a colonial choanoflagellate containing both flagellated and amoeboid cells appeared in 1882^29^, but was never corroborated^22^, raising questions about its validity. Instead, the diagnostic and seemingly universal cell architecture of choanoflagellates is that of a rigid, ovoid cell bearing an apical collar complex – a single flagellum surrounded by a microvillar collar. The close evolutionary relationship between animals and choanoflagellates^20^, coupled with the similarity of morula stage animal embryos to spherical colonies of choanoflagellates^30^ lent apparent support to Metschnikoff’s hypothesis^25^ and led us and others to infer that the amoeboid cell types of animals had evolved from ancestral flagellate cells^25,31–33^ after the establishment of multicellularity. However, modern choanoflagellates doubtless differ in some respects from their last common ancestor with animals^34^ and some close outgroups to choanoflagellates and animals produce amoeboid cells^3,8,20,35,36^ (some of which have recently been shown to alternate with a flagellate form^37–39^), raising the possibility that the cellular machinery for cell crawling and flagellar swimming both predate the divergence of the choanoflagellate and animal lineages^16^. Intriguingly, rare and transient episodes of cell deformation have been reported to precede cell division in certain types of choanoflagellates^22^. Moreover, the phenotypic repertoire of modern choanoflagellates is likely not completely known, as several fundamental aspects of choanoflagellate biology have only been discovered in the past few years, including collective contractility^40^, sexual reproduction^41^, and bacterial regulation of multicellularity^42^ and mating^43^.

We report here on our recent and serendipitous discovery of environmentally-relevant conditions under which the choanoflagellate *Salpingoeca rosetta* transdifferentiates from a flagellated state into an amoeboid state. While growing *S. rosetta* under conditions in which the cells were physically confined by medium evaporation, we observed that some cells transitioned from a flagellated state to an amoeboid state (Supplementary Video 1). Although amoeboid cells have not previously been reported in choanoflagellates, physical confinement regulates amoeboid cell differentiation and crawling motility in a wide range of eukaryotic cells, including zebrafish embryonic cells^9^, mammalian mesenchymal cells^11^, chytrid fungi^39^, dictyostelid amoebae^44^, and euglenoid algae^45^. Moreover, cell confinement is likely of ecological relevance for choanoflagellates, which have been detected in diverse granular microenvironments (soils^46^, marine sediments^47,48^, sands^49^, and silts^49^) whose pore sizes range from 1 mm to < 1 µm and extend below the range of typical choanoflagellate cell diameters (∼2 to 10 µm)^22^.

To test whether mechanical cues can induce the amoeboid phenotype, we used a tunable system for dynamic cell confinement^50^ and imaged live *S. rosetta* cells before, during, and after cell confinement (Fig. 1a). Single cells of *S. rosetta* confined in a space of 4 µm or more maintained the canonical flagellate phenotype, consistent with the cell body not being deformed. On the other hand, confinement below 3 µm elicited an active response from the cells, which started dynamically extending and retracting protrusions within a few seconds (Fig. 1b-d; Supplementary Video 2). In 2 µm confinement or less, most cells retracted their collar complex (the apical flagellum and microvillous collar) within minutes, thus acquiring a fully amoeboid phenotype (Fig. 1b-j; Extended Data Fig. 1a; Supplementary Video 2). Releasing confinement fully reversed the phenotypic switch (Fig. 1k-p). Newly unconfined cells retracted their dynamic protrusions, regained a round shape, and regrew a flagellum close to the position of the original, retracted one (Extended Data Fig. 1b-d; Supplementary Video 3), suggesting that information on apicobasal cell polarity is conserved in the amoeboid form, though not externally visible.

**Figure 1.**
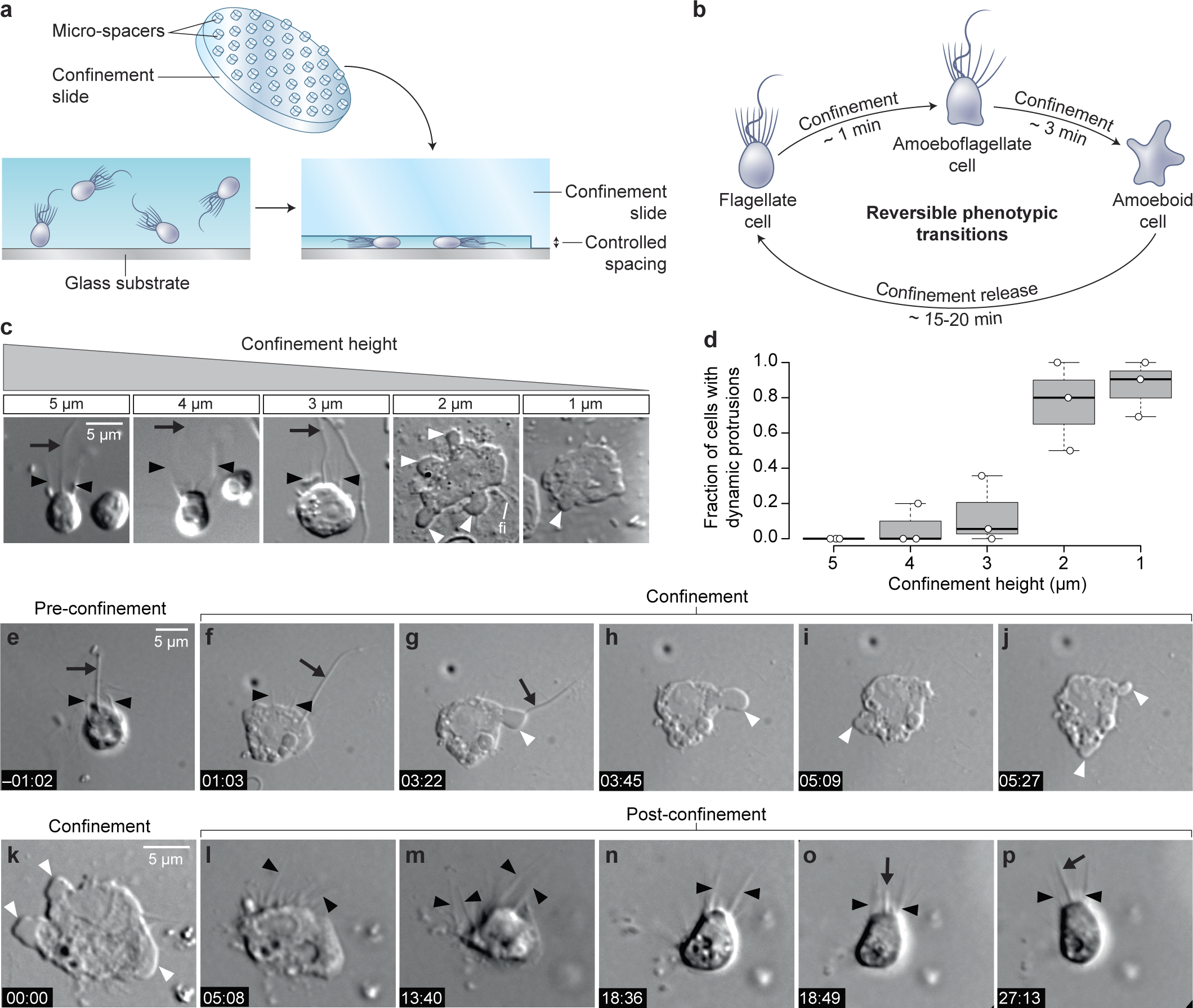
Confinement induces an amoeboid phenotype in the choanoflagellate *S. rosetta*. **a**, Free-swimming cells (bottom left) were confined (bottom right) at a fixed height using confinement slides with micro-spacers^11,50^ (top). **b**, Confined *S. rosetta* cells underwent a rapid phenotypic transition, first from a flagellate form into an amoeboflagellate form, and eventually into an amoeboid form. Releasing confinement reversed this transition. **c, d**, Confinement height correlated with the phenotypic switch. **c**, Representative cells at each confinement height tested. **d**, The flagellate form dominated at >3 µm confinement and the amoeboid form (defined by the presence of dynamic protrusions) at <3 µm. **e-j**, Time series of an *S. rosetta* cell switching to the amoeboid form at 2 µm confinement. See Supplementary Video 2 for multiple examples. **k-p**, Time series of an amoeboid *S. rosetta* cell reverting to the flagellate form after release from confinement. See Supplementary Video 3 for multiple examples. In all panels, white arrowheads indicate dynamic protrusions, black arrowheads indicate collar microvilli, and black arrows indicate the flagellum. Time stamps in black boxes shown as min:sec.

In addition to the solitary “slow swimmer” cells used in the experiments above, *S. rosetta* can also differentiate into other cell types, including multicellular rosettes and sessile “thecate” cells^30^. Both these cell types also differentiated into amoeboid cells with dynamic protrusions under confinement, showing that competence to undergo the amoeboid switch is not restricted to a single *S. rosetta* cell phenotype (Extended Data Fig. 2). Finally, *S. rosetta* responded in the same way to every type confined environment tested, including the pressure-controlled dynamic cell confiner, glass coverslips separated by microbeads (serving as spacers) (Supplementary Video 4), thin liquid films spread under a layer of oxygen-permeant oil (Supplementary Video 5), and agar gels (Supplementary Video 6). This suggests that the amoeboid switch is induced by cell deformation itself, independently of the properties of the substrate.

**Figure 2.**
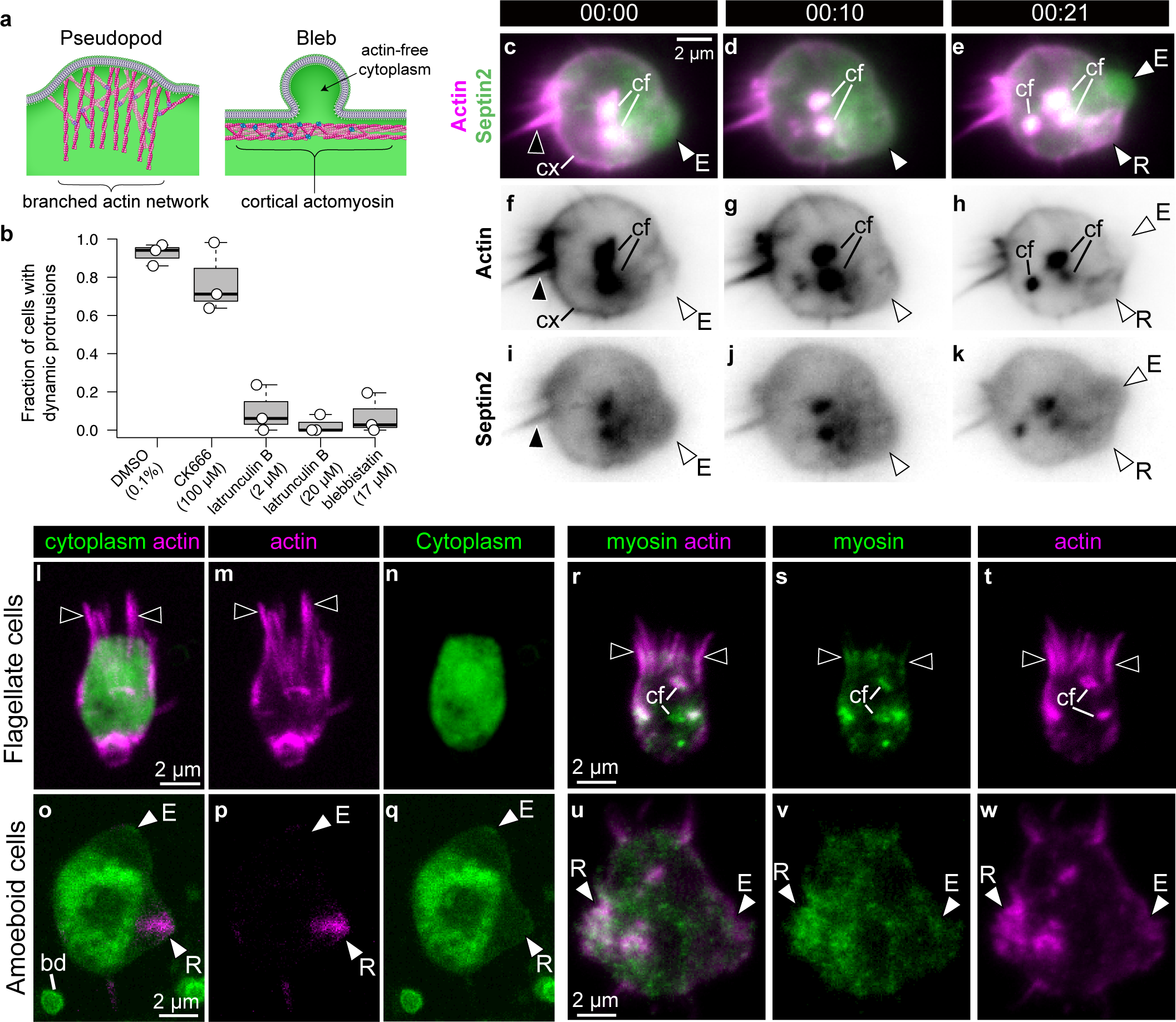
*S. rosetta* amoeboid cells generate blebs. **a**, Protrusions in eukaryotic crawling cells can either be F-actin-filled pseudopods that form by polymerization of F-actin (pink) reticulated by the Arp2/3 complex (purple, left), or F-actin-free blebs that form through the action of contractile forces in the actomyosin cortex underlying the plasma membrane (right). The cytosol is green in both panels. Modified from^52^. **b**, Formation of dynamic protrusions required F-actin and myosin II activity, but not Arp2/3-mediated F-actin polymerization. Protrusions were abundant in DMSO-treated control cells (N=116) and in cells treated with the Arp2/3 inhibitor CK666 (100 µM, N=162) but virtually absent in cells treated with the F-actin polymerization inhibitor latrunculin B (at both 2 µM (N=120) and 20 µM (N=103)) or the myosin II inhibitor blebbistatin (17 µM, N=130). This suggests that the dynamic protrusions were blebs. All cells were under 1 µm confinement. **c-k**, Dynamic protrusions that form under confinement are blebs, as indicated by live imaging of an *S. rosetta* amoeboid cell expressing an F-actin marker (LifeAct-mCherry, magenta) and a tagged septin (septin2-mTFP, green – which distributes throughout most of the cytoplasm when expressed at high levels). **c, f, i, and e, h, k**, Expanding blebs (E) were cytoplasm-filled but F-actin-free. **e, h, k**, Retracting blebs (R) were re-invaded by F-actin. F-actin was also present as a cortical layer (cx), as in animal cells^85^, and accumulated in cytoplasmic foci (cf). The cell was under 2 µm confinement. **l-q**, *S. rosetta* cells fixed and stained for F-actin (phalloidin, magenta) and cytoplasm (FM 1-43 FX, which distributes to the cytoplasm of *S. rosetta* following fixation, green) confirmed that the protrusions of amoeboid cells initially lack F-actin and are therefore blebs. **l-n**, A flagellate cell showing collar microvilli (black arrowheads). **o-q**, An amoeboid cell showing both F-actin-free and F-actin-filled protrusions, respectively, interpreted as expanding (E) and retracting (R) blebs. bd: 1 µm microbead used as confinement spacer. **r-t**, In *S. rosetta* flagellate cells fixed and stained for F-actin (phalloidin; magenta) and myosin II (CMII 23 antibody^40^; green), actin localized primarily to the microvillar collar and cytoplasmic foci. **u-w**, Notably, in amoeboid cells F-actin and myosin II colocalized in the presumptive retracting blebs (R), but not in the presumptive expanding blebs (E). In all panels: white arrowheads: blebs, black arrowheads: microvilli, E: expanding blebs, R: retracting blebs, cf: cytoplasmic foci. Time stamps in black boxes shown as min:sec.

To reconstruct the evolutionary history of a given cellular phenotype (such as the amoeboid phenotype), an important pre-requisite is the identification of the cellular and molecular modules that underlie it in a phylogenetically relevant set of species^8,51^. Eukaryotic cell protrusions similar to those we observed in *S. rosetta* fall into two different categories, pseudopods and blebs, that differ in their underlying mechanisms^52^. Pseudopods contain branched F-actin networks reticulated by the Arp2/3 complex and have been best-studied in adhesive animal mesenchymal cells^53^ but have also been identified in chytrid fungi^39^. By contrast, blebs form as actin-free protrusions by detachment of the plasma membrane under the influence of actomyosin cortex contractility^54^ (Fig. 2a) and have been well-documented in migratory primordial germ cells^17^. The presence of pseudopods and blebs is not mutually exclusive and multiple cells, such as animal mesenchymal cells^55^ or dictyoselid amoebae^56^, can produce both.

To determine the nature of the *S. rosetta* cell protrusions, we treated cells with small molecule inhibitors of proteins required for pseudopod or bleb formation in animals and other amoeboid lineages. Both pseudopods and blebs require the presence of F-actin networks. As expected, we found that inhibition of actin polymerization with latrunculin B prevented the formation of cell protrusions in *S. rosetta* (Fig. 2b). Inhibition of the Arp2/3 complex with CK666 did not prevent formation of *S. rosetta* cell protrusions, suggesting they might represent blebs rather than pseudopods (Fig. 2b). The inference that these protrusions are blebs was independently demonstrated by disruption of actomyosin contractility by treatment with the myosin II inhibitor blebbistatin, which entirely abolished formation of cell protrusions (Fig 2b).

In animal cells, expanding blebs are initially devoid of F-actin, which then re-invades the blebs before retraction^57^. Using live *S. rosetta* cells transfected with LifeAct-mCherry^58^, we observed similar F-actin dynamics in the protrusions of confined cells (Fig. 2c-k; Supplementary Video 7). Because the use of LifeAct can sometimes create artifacts^59^, we further studied F-actin localization in fixed confined cells stained with fluorescent phalloidin and a marker of the cytoplasm. Again, we observed both actin-free and actin-filled protrusions, consistent with our observations of expanding and retracting blebs, respectively (Fig. 2l-q; Extended Data Fig. 3).

**Figure 3.**
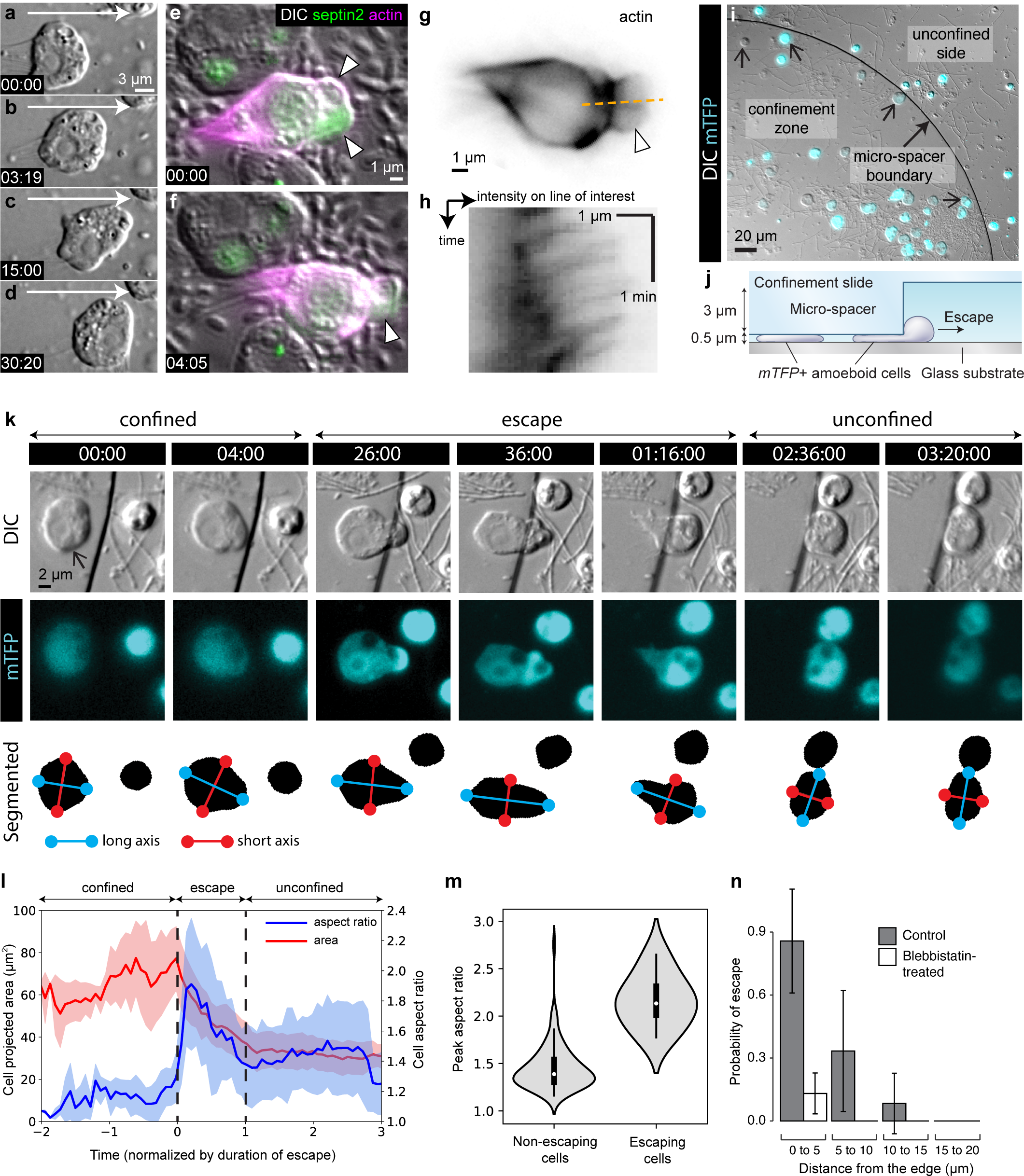
The amoeboid switch allows escape from confinement. **a-d**, Amoeboid cell crawling after flagellar retraction (Supplementary video 2). White arrow indicates direction of movement. **e-h**, Live imaging of a cell expressing LifeAct-mCherry (magenta) and a cytoplasmic marker (green) revealed the dynamics of the actin cytoskeleton during crawling (white arrowheads: blebs; Supplementary video 9). **e, f**, Two time points showing crawling behavior of a cell expressing LifeAct-mCherry and a cytoplasmic marker. **g**, Analysis of F-actin distribution along a line of interest (dotted orange line) in a crawling cell (LifeAct channel from panel **f** depicted in inverted intensity) over time (**h**) allowed visualization of actin flow. **h** is a kymograph showing retrograde flow over the line of interest in the LifeAct-mCherry-expressing cell. F-actin underwent retrograde flow inside the bleb, similar to animal amoeboid cells under confinement^9,11^. **i**, mTFP-expressing *S. rosetta* cells (cyan; confined cells that escaped during the assay indicated with small arrows) distributed within and outside the confinement zone (border indicated with larger arrow) at the beginning of an escape assay (Supplementary video 10). **j**, Schematic of cross-section through escape assay set-up from **i. k**, Time series of an mTFP-expressing cell (arrow) during escape from confinement (top, DIC; middle, mTFP; bottom, segmentation of mTFP fluorescence to reveal cell shape; Supplementary video 11). Automated detection of the long (blue) and short (red) axes of the cell revealed that the cell elongated during crossing of the confinement border and relaxed into a more rounded shape once escape was complete. **l**, Escaping cells (N=8) consistently elongated during escape and resumed a rounder shape once in the unconfined area. Escape also corresponded to a decrease in the projected area of the cell. Mean aspect ratio (red line) and projected area (blue line), ribbons: standard deviation. **m**, Escaping cells (N=8) acquired a highly elongated shape. Non-escaping cells (N=86) did not reach comparable elongation levels, as indicated by the peak aspect ratio (*p*=4.5*10^−6^ by Mann-Whitney’s U test). **n**, Escape required myosin II activity. Control cells (N=142) almost always escaped confinement if they were initially located less than 5 µm away from the border, and some escaped from as far as 15 µm. 17 µM blebbistatin-treated cells (N=183) virtually never escaped. Time stamps in black boxes shown as min:sec.

In animal cells, bleb formation is caused by the contractile activity of myosin II^54^ which co-localizes with actin in the cell cortex and within retracting blebs^57^. As myosin II inhibition disrupts bleb formation (Fig. 2b), we hypothesized that cortical actomyosin might also underlie blebbing in *S. rosetta*. Immunofluorescence confirmed the presence of actomyosin in the cortex of non-confined flagellate cells (as previously reported^40^; Fig. 2r-t) and in the retracting blebs of confined cells (Fig. 2u-w).

Most confined *S. rosetta* cells remained in one place and extended blebs in all directions without net locomotion. However, a few cells did migrate the cells over short distances (about 15 µm; Fig. 3a-f; Supplementary Video 1, 2, and 9). Like animal crawling cells^9,11,13^, the *S. rosetta* cells that migrated underwent marked retrograde F-actin flow (Fig. 3g-h).

One of the regulators of actomyosin activity in some animal cells^14,60^ and in *Dictyostelium*^61^ is intracellular microtubule distribution, with microtubule-free zones experiencing higher local contractility and bleb retraction. In unconfined flagellated *S. rosetta* cells, we observed that cortical microtubules radiated from the apical basal body to form a cage underneath the entire plasma membrane^62,63^ (Extended Data Fig. 4 a-d). In amoeboid cells, this cage remained present – but mostly detached from the plasma membrane and around the nucleus (Extended Data Fig. 4 e-h). This is consistent with maintenance of the microtubule-organizing center and of apicobasal polarity in amoeboid cells (Extended Data Fig. 1). Interestingly, microtubule depolymerization induces blebbing in *S. rosetta* in the absence of confinement (Extended Data Fig. 4 i-p; Supplementary Video 8). This suggests that, as in animal cells, microtubule distribution might regulate blebbing.

**Figure 4.**
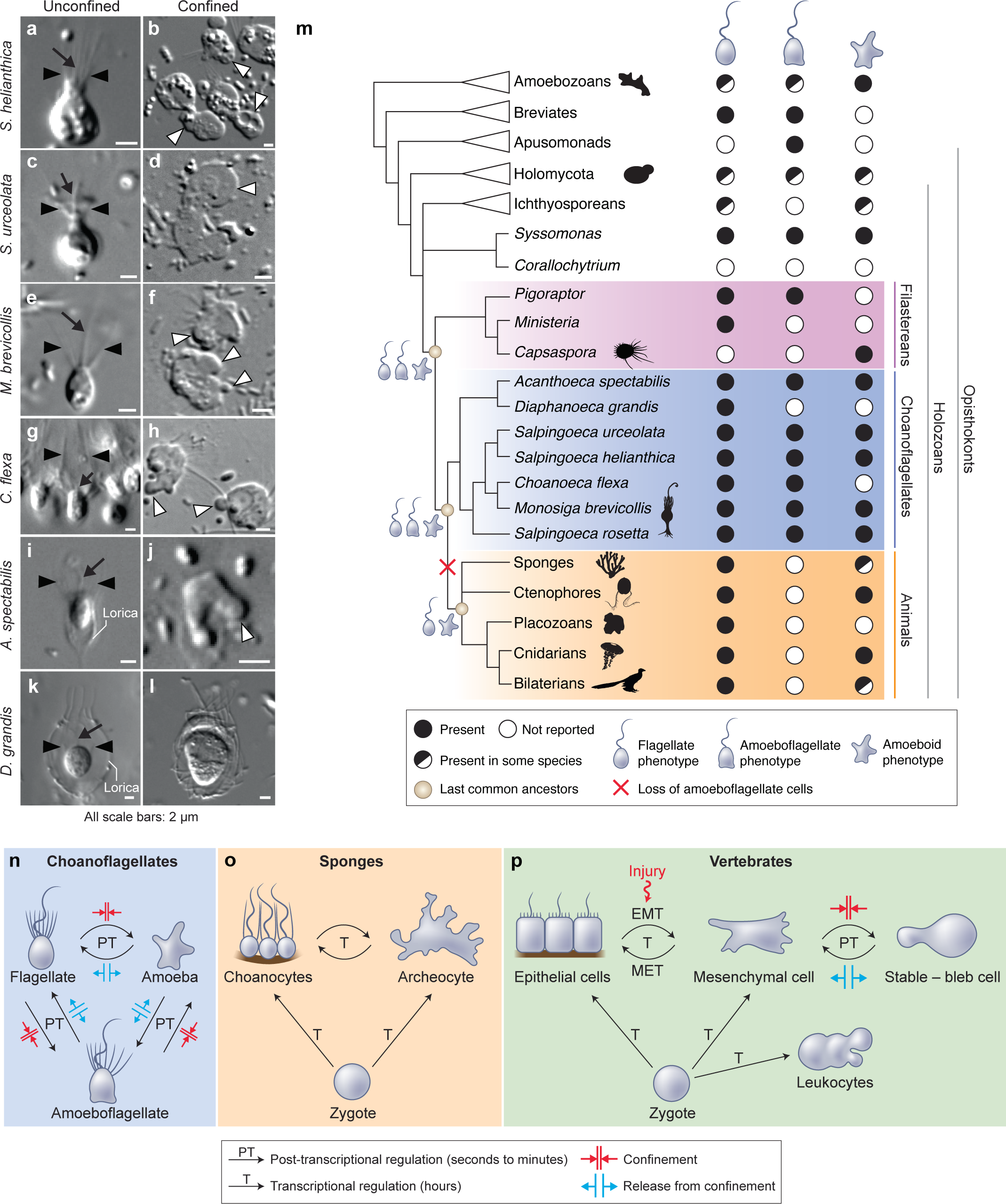
The last common choanozoan ancestor had amoeboid and flagellate life history stages. **a-j**, Five of six choanoflagellate species tested underwent the amoeboid transition under 2 µm confinement (Supplementary Videos 14-17). **k-l**, In contrast, the loricate choanoflagellate *Diaphanoeca grandis* was passively flattened under 2 µm confinement, but did not generate blebs (Supplementary Video 13). **m**, Phylogenetic distribution of flagellate, amoeboflagellate, and amoeboid cell phenotypes in animals, fungi, amoebozoans, and their relatives. We infer that the last common ancestor of choanoflagellates and animals was able to differentiate into flagellate, amoeboid, and amoeboflagellate forms. Flagellate and amoeboid forms were likely both still present in the last common ancestor of all animals. See Expanded Data Figure 6 and Supplementary Table 1 for full supporting evidence regarding the distribution of cellular phenotypes in animals. Species silhouettes are from Phylopic (http://phylopic.org). **n-p**, Commonalities and differences in the regulation of cellular phenotypic transitions in choanoflagellates, sponges, and vertebrates. **n**, Choanoflagellates can rapidly alternate (in a matter of minutes) between flagellate, amoeboflagellate, and amoeboid forms based on degree of external confinement. Inhibition of transcription does not prevent transitions between amoeboid and flagellate forms, suggesting that the transition is post-transcriptionally regulated (Extended Data Fig. 8). **o**, In sponges, the zygote can give rise to flagellated choanocytes and amoeboid archeocytes. Choanocytes and archeocytes can reversibly interconvert, but this process takes several hours and likely requires transcriptional regulation^34^. **p**, In vertebrates, multicellular development from the zygote results in terminal differentiation of ciliated epithelial cells, mesenchymal cells and amoeboid leukocytes, but injury can trigger differentiation of epithelial cells into crawling mesenchymal cells (by epithelial-to-mesenchymal transition, or EMT) that are respond to confinement switching into a stable bleb form that is capable of amoeboid migration^11^. The switch from epithelial to mesenchymal cells is reversible (by mesenchymal-to-epithelial transition, or MET).

These results show that in choanoflagellates – as in animal cells and slime molds – confinement induces blebbing, which relies on actomyosin activity and appears regulated by microtubule distribution. One difference, however, is that calcium signaling does not appear to be required for confinement-induced blebbing in *S. rosetta* (Extended Data Fig. 5).

The amoeboid switch (at least as induced through confinement between glass slides) resulted in cells that extendeded blebs and seemed to probe their local environment, but only rarely migrated. Thus, we hypothesized that it might enable an escape response in a more complex environment, closer to natural interstitial media. To test this hypothesis, we tracked the behavior of live cells in a heterogeneous environment containing zones of confinement surrounded by less confined spaces (Fig. 3i-j). A matrix of PDMS pillars defined an environment in which choanoflagellates encountered 0.5 µm-deep confinement zones surrounded by 3.5 µm-deep spaces in which they could swim freely. Cells that were initially confined less than 15 µm away from a pillar border were capable of escaping to a 3.5 µm-deep space (Fig. 3k; Supplementary Video 10). Typically, a confined cell would first bleb irregularly and crawl slowly, until part of the cell – generally an expanding bleb – crossed the border of the pillar into the non-confined space. Following this, the cell would change shape and elongate away from the border of the pillar and crawl directionally until it had fully escaped (Fig. 3k; Supplementary Video 11). Automated cell segmentation followed by morphometric quantification confirmed that escaping cells reliably elongated (Fig. 3l; Extended Data Fig. 6; Supplementary Video 11) while non-escaping cells (that never detected a border) remained nearly round (Fig. 3m), indicating that this escape behavior involves specific cell shape changes. Interestingly, the first blebs that crossed the border were occasionally shed from the cell and reabsorbed during escape (Supplementary Video 12). Finally, treatment with blebbistatin dramatically reduced the escape behavior (Fig. 3n), consistent with this behavior requiring myosin activity. This type of behavior might allow choanoflagellates to escape from tightly packed silts (< 3 µm granularity) into the water column or more loosely packed interstitial environments.

Having uncovered an amoeboid switch in *S. rosetta*, we sought to assess the phylogenetic distribution of the amoeboid switch across choanoflagellate diversity in order to determine whether this phenotype may have been present in the last common ancestor of choanoflagellates. We tested the effect of 2 µm confinement on six additional choanoflagellate species that together cover all main branches of the choanoflagellate phylogenetic tree^64^. All displayed blebbing activity under confinement (Fig. 4a-j) with the exception of *Diaphanoeca grandis* (Fig 4k,l; Supplementary Video 13) – which could indicate a secondary loss of the ameboid switch in this lineage, or that different conditions are required to induce the phenotype (such as <2 µm confinement). The amoeboid switch in the other five species covers a spectrum of phenotypes. The least pronounced response is seen in *Choanoeca flexa* (Fig. 4g,h), whose sheet colonies^40^ dissociate into single cells that bleb and migrate over short distances without retracting their flagella, thus keeping an amoeboflagellate phenotype. Two other species, *Monosiga brevicollis* (Fig. 4e,f; Supplementary Video 14) and *Acanthoeca spectabilis* (Fig. 4i,j; Supplementary Video 15) showed a similar response to *S. rosetta*, including blebs and flagellar retraction. *Salpingoeca helianthica* generated very large and branched blebs (Fig. 4a,b; Supplementary Video 16), often longer than the rest of the cell body – reminiscent of the “lobopods” described in some protists^38^. Finally, *Salpingoeca urceolata* (Fig. 4c,d; Supplementary Video 17) differentiated into amoebae capable of sustained migration over long distances (>40 µm).

The discovery of an amoeboid switch in a choanoflagellate has the potential to illuminate the ancestry of amoeboid cells in animals. Crawling cell motility is found in virtually all animal lineages and is important for embryonic development (e.g. in neural crest migration^65^, gastrulation^66^ and primordial germ cell migration^17,18,67^), wound healing^19^, and immunity (e.g. in phagocytes patrolling tissues^14^). Cell crawling has also been frequently observed in the sister-lineage of choanozoans, filastereans^36–38^ – but its seeming absence from choanoflagellates had made its evolutionary history unclear. Together with the existing comparative evidence, our data suggest that the last common ancestor of choanoflagellates and animals had the ability to differentiate into amoeboid and amoeboflagellate cells under confinement.

Consistent with an ancient origin of all three phenotypes, amoeboid, flagellate and amoeboflagellate cells are all broadly distributed in opisthokonts (animals, fungi, and their relatives). Interestingly, amoeboflagellate phenotypes have recently been described in several species occupying key phylogenetic positions, including in the sister-group of choanozoans (the filastereans)^37,38^, in early-branching fungi^39,68,69^, in the two closest known relatives of opisthokonts (apusomonads^70^ and breviates^71^), and in early-branching amoebozoans^72^. This is consistent with an ancient origin and broad conservation of the amoeboflagellate phenotype in opisthokonts^73^. Opisthokont ancestors might have crawled using a combination of blebs (as in amoeboid choanoflagellates, amoeboid animal cells^9,11^, and amoebozoans^44^), pseudopods (which are present in choanoflagellates during phagocytosis^22,39^ and during crawling in chytrid fungi^8,39^, amoebozoans^44^, and some holozoans^38^), and filopodia (which contribute to locomotion in filastereans^63^ and choanoflagellate settlement^30^). Moreover, cell crawling is regulated by confinement in animal cells^9–12^, choanoflagellates, chytrid fungi^39^ and dictyostelid amoebozoans^44^, suggesting that the ability to respond to confinement might be an ancient feature. These prior findings, together with our observation of an amoeboid switch in choanoflagellates, suggest that the switch from a flagellate to a crawling phenotype in response to confinement was part of an ancestral stress response in the last common choanozoan ancestor^74^. The crawling behavior of the choanozoan ancestor might even have been more extensive than what we observed in modern choanoflagellates: indeed, most choanoflagellate species have secondarily lost some genes often involved in crawling motility, such as the integrin complex^75,76^ and the transcription factors Brachyury^77^ and Runx^1,75^. Interestingly, the only choanoflagellate species known to have retained Runx is *S. urceolata*^75^, which displayed the most extensive crawling behavior under confinement (Fig. 4d, Supplementary Video 17). Another intriguing observation lies in the fact that animal cells with artificially disrupted components of the integrin adhesome show little motility between flat surfaces but retain the ability to crawl through constrictions^13^ – reminiscent of our observation of the *S. rosetta* escape response.

Specialized crawling cell types are present in multiple animal lineages, including sponges (archeocytes), ctenophores (stellate cells), cnidarians (amoebocytes), invertebrate bilaterians (ameobocytes) and vertebrates (white blood cells and mesenchymal cells) (Supplementary Table 1; Extended Data Fig. 7). We propose that these might have evolved by stabilization of an ancestral stress response to confinement, representing an example of evolution of a cell type from temporally alternating phenotypes (in line with the “temporal-to-spatial transition” hypothesis^1,3,15^) and more specifically from a stress response^78^. During this transition, the switch between a flagellate phenotype and a crawling phenotype – ancestrally fast and post-transcriptionally regulated, as in modern choanoflagellates (Fig. 4n; Extended Data Fig. 8) – would have come under the control of transcriptional regulators^19^ (Fig. 4o,p). This switch would have remained reversible at first (as it still is in sponges^34^; Fig. 4o) but later become irreversible with the evolution of terminal cell differentiation (Fig. 4p). However, the ancient response to confinement was not necessarily fully lost – as confinement activates blebbing in vertebrate mesenchymal and embryonic cells^9,11^ through a transduction pathway that makes use of newly evolved proteins, such as the phospholipase cPLA2^10,12^ that does not exist in choanoflagellates^21,75^.

Although constitutively crawling cell types are widespread in animals, they seem absent from calcaronean sponges^79^, placozoans^80^, and xenacoelomorph worms^81^. In these lineages, cell crawling often exists instead as a transient phenomenon. Indeed, in many animal lineages (including those with stable amoeboid cell types), transient cell crawling often contributes to embryonic development (for example, primordial germ cells often display amoeboid migration^18,67^; Supplementary Table 1) and/or to wound healing (which often involves crawling by cell types that do not normally display it, such as epithelial cells^19^; Supplementary Table 1; Extended Data Fig. 7). Further research will be required to determine how many times this transient cell behavior was converted into a stable cell type during animal evolution. This conversion could have happened once (in a common ancestor to all animals) or multiple times independently in several lineages. Another open question concerns the nature of the mechanotransduction pathway by which choanoflagellates detect and respond to confinement, although it might involve microtubule detachment from the plasma membrane (Extended Data Fig. 4). Finally, future comparative work will benefit from deeper insights into the mechanisms of cell crawling in choanoflagellates and animals^8^. Intriguingly, the amoeboid cell types of sponges (archeocytes) express more genes shared with choanoflagellates than other sponge cell types do^34^ (including choanocytes), which might include genes involved in crawling motility^3^. Functional characterization^82^ of those genes^8^ in multiple phylogenetically relevant species as well as large-scale efforts to map animal cell type diversity within a phylogenetic framework^83,84^ will help reveal how cell types have evolved and diversified during animal evolution.

## Methods

### Choanoflagellate cultures

#### *Cultures of* S. rosetta

*Salpingoeca rosetta* in the chain/slow swimmer form^30^ was maintained as a co-culture with the prey bacterium *Echinicola pacifica* (SrEpac) in 5% Sea Water Complete (SWC) culture medium, as previously described^41^. Thecate *S. rosetta* were from a thecate SrEpac strain (HD1) which was produced from SrEpac through starvation following a published protocol^41^. Rosettes were obtained from a co-culture of *S. rosetta* with the multicellularity-inducing bacterium *Algoriphagus machipongonensis*^30,42^ (strain Px1) in 5% Cereal Grass Medium (CGM3) in Artificial Sea Water (ASW)^86^.

#### *Cultures of* C. flexa, S. helianthica, S. urceolata, D. grandis, M. brevicollis, *and* A. spectabilis

Cultures of *S. urceolata, D. grandis, M. brevicollis* and *A. spectabilis* were established by thawing frozen stocks stored in liquid nitrogen following a published protocol^87^. Recipes for the culture media were previously published^75,86^ and modified as follows: *S. urceolata* was grown in 1% CGM3 at 25°C, *D. grandis* was grown in 5% CGM3 at 16°C and *A. spectabilis* was grown at 16°C. *C. flexa* was obtained from a culture founded by a colony isolated from Curaçao in 2018 and continuously passaged since then as previously published^40^. Live cultures of *S. helianthica* were a gift from Mimi Koehl and Michael O’Toole II and were maintained in 25% freshwater CGM3 (FCGM3) following a published protocol^75^.

#### Live imaging

Cells were imaged by differential interference contrast (DIC) or epifluorescence microscopy using a 40x (water immersion, C-Apochromat, 1.1 NA), 63x (oil immersion, Plan-Apochromat, 1.4 NA), or 100x (oil immersion, Plan-Apochromat, 1.4 NA) Zeiss objective mounted on a Zeiss Observer Z.1 with a Hamamatsu Orca Flash 4.0 V2 CMOS camera (C11440-22CU).

#### Confocal imaging

Fixed and stained samples were imaged by confocal microscopy using a Zeiss LSM 880 AxioExaminer with Airyscan and a 63x, 1.4 NA C Apo oil immersion objective (Zeiss) and excitation provided by a 405, 488, 568, or 633 nm laser (Zeiss).

### Cell confinement

#### Dynamic cell confiner

A 1-well dynamic cell confiner^11,50^ comprising an Elveflow Vacuum/Pressure Generator and an Elveflow AF1 DUAL–Vacuum/Pressure Controller was purchased from 4Dcell (Montreuil, France) together with suction cups and with 1 µm, 2 µm, 3 µm, 4 µm, and 5 µm confinement slides.

All confinement assays were realized on cells mounted in FluoroDishes (World Precision Instruments FD35-100) under a confinement slide and imaged on a Zeiss Observer Z.1 (see above). *S. rosetta* dynamic confinement experiments were performed using SrEpac cultures that were dense, but not starving (∼10^6^ cells/mL). In assays aiming at visualizing both flagellar retraction and regeneration in the same cells (Extended Data Fig. 1, Supplementary Video 3), the cells were attached to the substrate by coating the FluoroDish with 0.1 mg/mL poly-D-lysine for 1 minute (Sigma Aldrich P6407-5MG; washed twice quickly with ASW) before mounting the cells. In assays aimed at investigating the possibility of crawling (Supplementary Video 2 and escape assays), poly-D-lysine was omitted.

Confinement was applied by following provider’s instructions, by gradually decreasing pressure from −3 kPa to −10 kPa with the vacuum/pressure controller. Confinement was released by gradually restoring pressure to −3 kPa.

Escape assays were realized following the same protocol as confinement assays, but by imaging the cells trapped under the micropillars of a 3 µm confinement slide.

#### Confinement with microbeads

Some early confinement experiments and pharmacological assays were performed by confining *S. rosetta* cell suspensions between two coverslips separated by microbeads (acting as spacers) and imaging them on a Zeiss Z.1 observer (see above).

The following types of microbeads were used: non-fluorescent 1 µm sulfate/latex beads (ThermoFisher Scientific S37498), non-fluorescent 2 µm sulfate/latex beads (ThermoFisher Scientific S37500), orange fluorescent 2 µm microbeads (Sigma Aldrich L9529-1ML) and orange fluorescent 1 µm microbeads (Sigma Aldrich L9654-1ML). Beads were stored at 4°C and resuspended in ASW prior to experimentation by centrifugation for 10 minutes at 10,000 g on a tabletop microcentrifuge followed by supernatant removal and resuspension.

Prior to confinement, 100 mL of a dense *S. rosetta* culture (strain SrEpac, ∼10^6^ cells/mL) was filtered through a 5 µm syringe-top filter (Fisher Scientific SLSV025LS; to remove large biofilm pieces) and Percoll-purified (to remove bacteria) following a published protocol^41^. The resulting *S. rosetta* suspension was further concentrated into 100 µL by centrifugation at 5,000 g for 5 minutes on a tabletop microcentrifuge, thus reaching a final density of ∼10^9^ cells/mL. The resulting dense cell suspension was placed on ice and immediately mixed 10:1 with a stock suspension of microbeads in ASW. 0.1 µL of the cells/beads mixture was mounted on a rectangular coverslip pre-treated with a Corona surface treater (Electro-Technic Products BD-20AC) to facilitate liquid spreading, and surmounted with a second (non-Corona-treated) coverslip.

#### Confinement in a thinly spread liquid layer

The first confinement experiments (Supplementary Video 5) were realizing by trapping cells into a thinly spread layer of liquid medium surmounted by oxygen-permeant oil. 5 µL of a dense cell suspension (concentrated down to ∼10^9^ cells/mL in ASW complemented with 1% CGM3 and 1% rhodamine-dextran as a fluorescent marker of the aqueous phase (Sigma-Aldrich D6001)) were spread on a FluoroDish pre-treated with a handheld Corona surface treater and surmounted with 120 µL of oxygen-permeant anti-evaporation oil (Ibidi 50051). The thickness of the medium layer was measured by visualizing rhodamine-dextran fluorescence using a confocal microscope. Rhodamine-dextran fluorescence was exclusively observed within the aqueous layer of ASW-based medium (containing the cells) and was excluded from the overlaying oil (consistent with rhodamine-dextran being hydrophilic). Confocal stacks were visualized with Fiji^88^ and the decrease of red fluorescence at the water/oil interface allowed quantification of the thickness of the aqueous phase. Film thickness varied within a given FluoroDish (possibly due to meniscus effects) and ranged from 1 to 8 µm. Cells were observed to be consistently amoeboid if they were trapped in a layer thinner than 3 µm, and to be consistently flagellate and free-swimming if the layer was thicker than 5 µm – consistent with observations made with confinement slides and microbeads.

### Transfection

For F-actin live imaging (Fig. 2c-k), cells were co-transfected with plasmids encoding LifeAct-mCherry (Addgene ID NK612) and septin2-mTFP (Addgene ID NK641, which distributes within the entire cytoplasm in highly-expressing cells) following a published transfection protocol^58^ and imaged in epifluorescence microscopy using a Zeiss Z.1 observer.

Stable mTFP-expressing *S. rosetta* were produced following a previously published protocol^89^ by transfection^58^ of a plasmid encoding a puromycin resistance protein followed by mTFP, separated by a P2A self-cleaving peptide (Addgene ID NK676).

### Cell segmentation and morphometrics

For escape response assays (Figure 3), cell shapes were segmented using the DIC channel. Cell segmentations were obtained from the predictions of a StarDist model^90,91^. Ground truth for training the StarDist model was created by cropping out and manually labelling a subset of the cells to be analyzed, at evenly distributed timepoints throughout the movies. A total of N=160 square images (width 151px) of individual cells and their associated masks were rearranged into 32 mosaic images containing 5×5 cell images, representing the ground truth for network training. The centers of the segmented cells were tracked over time using Trackpy^92^.

Two distinct movies were analyzed, which led to 94 cells being segmented, of which 8 escaped. Cell centroid, aspect ratio, projected surface area, and circularity were computed from the segmented shapes. The confinement boundary was manually drawn as a circle at the position of the pillar border, which allowed computation of the distance between the boundary and the cell centroid, the cell front (defined as the minimal distance between the border and all points within the cell), and the cell rear (defined as the maximal distance between the border and all points within the cell).

For calcium depletion assays (Extended Data Fig. 5), as no strong difference in blebbing activity was readily observable between the different conditions, we set up a pipeline for automated quantification of blebbing activity to test for possible small, quantitative differences. The analysis was performed by M.A. who was blinded to the treatment conditions. Cells were segmented using the DIC channel. A StarDist model was trained using 14 manually labelled movie crops as ground truth (size 1024×1024px), containing comparable amounts of cells for each of the four different perturbations to be analyzed. Prior to prediction, the image background was estimated by applying a gaussian filter (kernel size 20px) and substracted from the input images. To improve the accuracy of the cell boundary reconstructions, the StarDist output was further processed by using the centers of the star-convex object predictions as seeds for a watershed segmentation based on the thresholded pixel predictions produced by the StarDist model’s UNet^93^.

Blebbing activity was approximated as the rate of cell shape change. First, all movies were resampled to a framerate of 1 frame per 20 s and only the first 240 s were considered. To compensate for a possible global drift of the field of view, all resulting frames of each movie were first registered to the first frame. For each cell and for each time point, the tracked cell labels were used to extract the difference between the shape of the cell and the shape of the same cell 20 seconds earlier. The zones over which cell shape differed between both time points matched recognizable blebs (see Supplementary Video 18 for an example). Blebbing activity for an individual cell was calculated as the mean area of this shape difference, averaged over all time points and normalized by cell area. Cell tracks shorter than 200s and those meeting any of the following conditions for any considered time point were excluded from this analysis:

- Cell area < 500px (pixel spacing: 0.1625 µm)
- Cell displacement between time points > 5px
- Cell area change between time points > 10 %
- Cell circularity < 0.7

Cell segmentation, tracking and all downstream analysis was performed in Python (3.7) in combination with software belonging to the SciPy ecosystem^94–97^ and additional software (CziFile: https://pypi.org/project/czifile/ and Fiji^88^). In the end, blebbing activity did not differ significantly between the four conditions tested (Extended Data Figure 5).

### Pharmacological assays

For all pharmacological inhibition assays, cells were pre-treated with small molecule inhibitors for 30 minutes before imaging (except for MBC, for which the pre-treatment was 36 hours). Negative controls were treated for the same time with a concentration of compound vector (most often DMSO) equivalent to that used in the highest inhibitor dosage. Compounds used and their stock and working concentrations are in the table below.

**Method Table 1.** Compounds used in pharmacological assays

| Methods Table 1. Compounds used in pharmacological assays |  |  |  |
| --- | --- | --- | --- |
| Name (target) | Provider | Stock solution | Working concentration |
| Blebbistatin (myosin II) | Sigma-Aldrich B0560 | 17 mM in 90% DMSO/10% H<sub>2</sub>O | 17 μM |
| Latrunculin B (F-actin) | Sigma-Aldrich L5288 | 10 mM in DMSO | 2-20 μM |
| MBC/Carbendazim (microtubules) | Sigma-Aldrich 378674 | 5 mg/mL in DMSO | 5 μg/mL |
| CK-666 (Arp2/3) | Sigma-Aldrich SML0006 | 100 mM in DMSO | 10-100 μM |
| BAPTA-AM (intracellular Ca<sup>2+</sup>) | ThermoFisher Scientific B6769 | 32.7 mM in DMSO | 327 μM |
| Actinomycin D | Sigma-Aldrich A1410 | 20 mg/mL in DMSO | 0.1 mg/mL |

### Calcium deprivation assays

For depletion of external calcium, cells were transferred into calcium-free AK sea water (CF-AKSW). CF-AKSW was prepared following a published AKSW recipe^58^, omitting only CaCl_2_, and further adding 20 mM EGTA to chelate any remaining calcium. A suspension of ∼10^8^ *S. rosetta* cells was concentrated into 100 µL (see “Cell confinement – Confinement with microbeads” section above) and resuspended in 1 mL CF-ASKW. Cells were then washed 3 times in 1 mL CF-AKSW in 1.5 mL plastic tubes by centrifugation (2 × 5 minutes and 1×15 minutes) at 10,000 g in a tabletop microcentrifuge. Cells were then confined using 2 µm microbeads as spacers as detailed above. Microbeads were similarly resuspended in CF-AKSW before being added to the cells.

For depletion of intracellular calcium, cells were incubated with 327 µM BAPTA-AM (a cell-permeant calcium chelator) for 30 minutes before confinement and imaging.

### Immunostainings of flagellate and amoeboid cells

Immunostainings of flagellated *S. rosetta* cells were performed following a previously published protocol^58^. Immunostainings of confined cells were performed by mounting cells between a small square coverslip (18×18 mm, VWR 470019-002) and a larger rectangular coverslip (24×50 mm, VWR 48393-241) using 1 µm or 2 µm microbeads as spacers (see “Cell confinement – Confinement with microbeads” above.) Immediately after confinement, the two lateral sides of the small coverslip (parallel to the long side of the large rectangular coverslip) were glued to those of the large coverslip using a small quantity of Super Glue gently spread with a micropipette tip (Methods Figure 1). This maintained close apposition of the two coverslips (and thus cell confinement) during the following steps. The Super Glue was left to dry for 5 minutes. This defined a flow chamber in which fixation, staining, and washing solutions could be pipetted on top of the large coverslip, close to the non-glued edges of the small coverslip, and then spread in the confined space by capillary action (Methods Figure 1).

**Method Figure 1.**
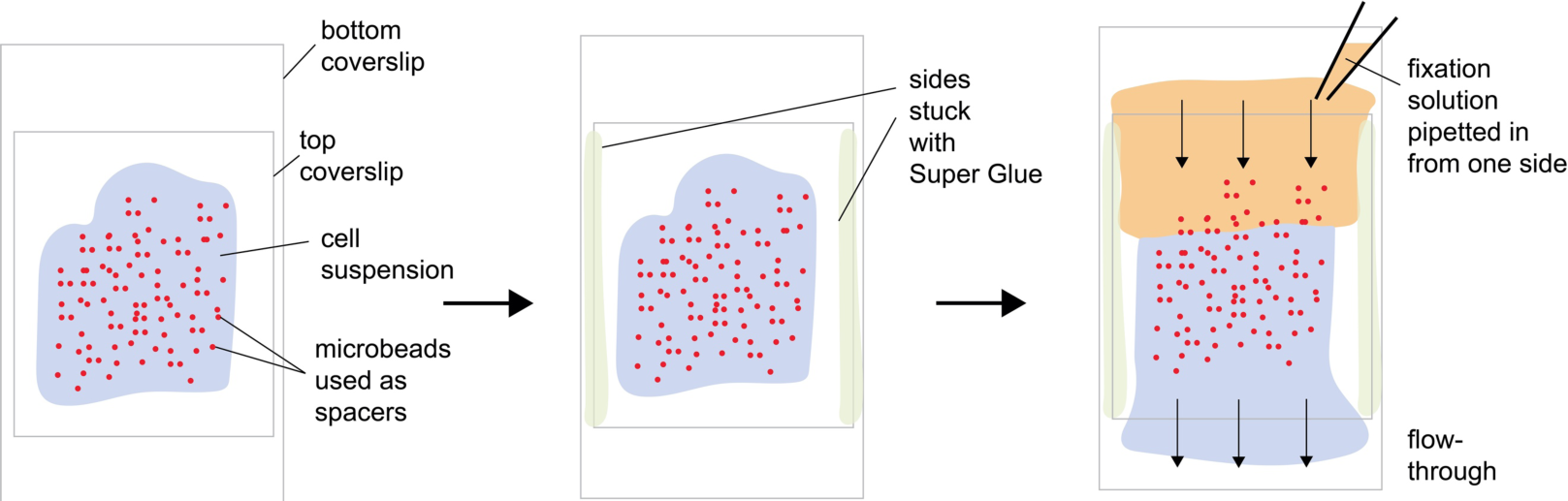
Flow-through chamber for immunostaining of confined cells.

50 µL of fixation solution (4% PFA in cytoskeleton buffer^58^) were pipetted close to the inflow side of the flow chamber (top in the last panel of Methods Figure 1). The slide was transferred into a small parafilm-sealed box, in which air humidity was maintained by water-soaked paper towels (to prevent evaporation of liquid in the sample). Fixation was left to proceed for 2 hours at room temperature. After each incubation step, excess liquid was removed by absorbing it with a Kimwipe (Fisher Scientific 06-666) on the outflow side (bottom in the last panel of Methods Figure 1) and the following solution was immediately added to the inflow side. After fixation, all washing steps were performed as when immunostaining flagellated cells, with the single difference that incubation in the primary antibody solution was allowed to proceed overnight at 4°C.

When included, 5 µg/mL FM 1-43 FX (Thermo Fisher Scientific F35355, dissolved in water as single-use aliquots) was mixed with the fixation solution. Primary antibodies were: mouse anti-β-tubulin (Developmental Studies Hybridoma Bank E7, diluted to 5 µg/mL – in Extended Data Fig. 4i); mouse anti-myosin II (Developmental Studies Hybridoma Bank CMII 23, diluted to 5 µg/mL); and rat anti-α-tubulin (YOL3/4, Abcam ab6161 diluted 1:300 – in Extented Data Figure 4a-h). Secondary antibodies were Alexa 488-anti-mouse (Invitrogen A11029) and Alexa 647-anti-rat (Thermo Fisher Scientific A-21247) diluted to the providers’ specifications. F-actin was stained with 0.66 units/mL Alexa 647-phalloidin (Life Technologies A22287) when combined with FM 1-43 FX, or 0.66 units/mL rhodamine-phalloidin (Life Technologies R415) when combined with myosin II immunostaining. DNA was stained with 10 µg/mL Hoechst 3342 (Thermo Fisher Scientific H3570).

## Acknowledgements

We thank D. Booth, F. Leon, M. Coyle, M. Sigg, and R. Aldayafleh for help with *S. rosetta* transfections; the staff and students of the 2018 Physiology course in the Marine Laboratory (Woods Hole, MA); O. Dudin for advice on MBC treatment; T. Linden, M. Coyle, D. Booth and F. Rutaganira for feedback on the manuscript; V. Ruprecht and L. Fritz-Laylin for feedback on the project; D. Maizels for assistance with the figures; and members of the King lab for stimulating discussions. T.B. was supported by the EMBO long-term fellowship (ALTF 1474-2016) and by the Human Frontier Science Program long-term fellowship (000053/2017-L).

## Author contributions

T. B. and N. K. designed the experiments and wrote the manuscript. T. B. performed most experiments. M. A. produced the automated cell segmentation and performed downstream image analyses. W. R. did early dynamic confinement experiments and imaged LifeAct-mCherry-transfected cells during the 2018 MBL Physiology course in Woods Hole. D. C. S. first observed the amoeboid switch in *S. helianthica*.

