## Supplementary Figures and Table for "A flagellate-to-amoeboid switch in the closest living relatives of animals"

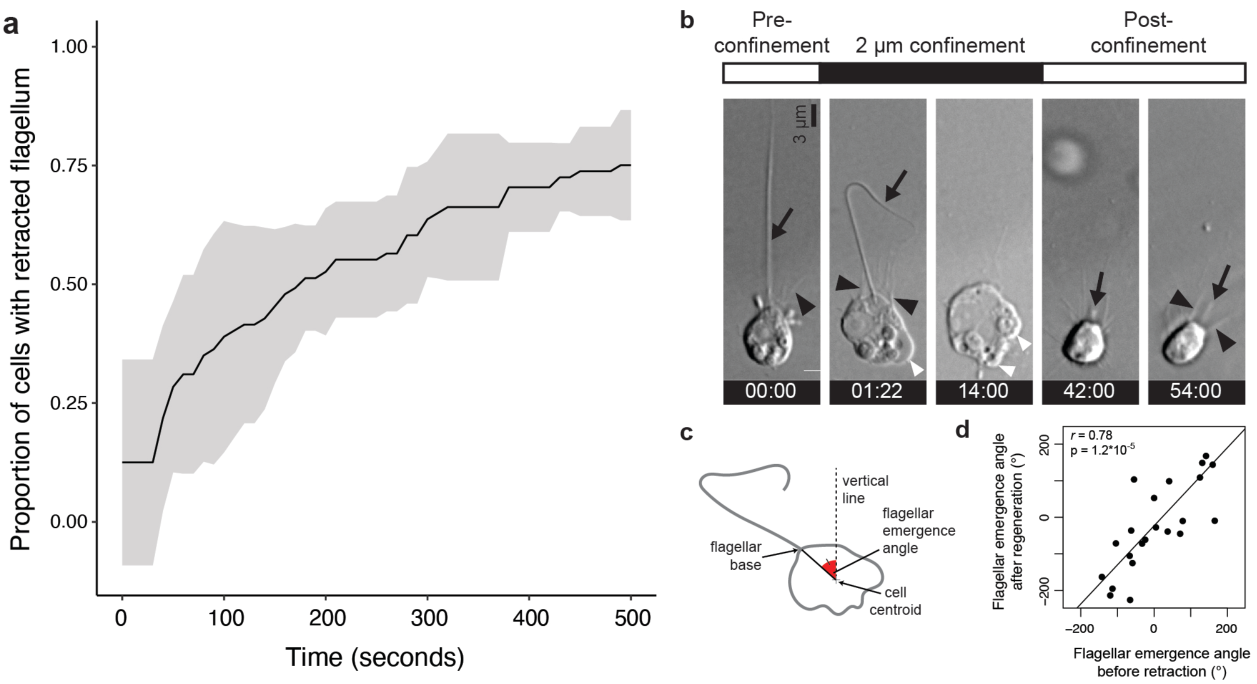


**Extended Data Figure 1. Flagellar retraction and regeneration during transitions between the flagellate and amoeboid forms**. **a,** Most (but not all) *S. rosetta* cells retracted their flagellum within 500 s of confinement at 2 µm. **b-d**, The flagellum regenerated after release from confinement in approximately the same position as the original, retracted flagellum. **b**, Time series of an *S. rosetta* cell retracting its flagellum under confinement and regenerating it after confinement release (Supplementary Video 3). The cell was attached to the glass substrate with poly-D-lysine to minimize cell movements. White arrowheads: dynamic protrusions, black arrowheads: microvilli, black arrow: flagellum. Time stamps in black boxes are min:sec. (Supplementary Video 3). **c,** To compare the flagellar position before and after confinement, the flagellar emergence angle was measured relative to an invariant vertical line (parallel to the edge of the field of view). **d**, The position of the regenerated flagellum after release from confinement in a population of cells was almost always close to the position of the original, retracted flagellum (as measured by the flagellar emergence angle; Supplementary Video 3). The cells sometimes underwent slight global reorientations under confinement (even though they were stuck to the substrate with poly-D-lysine) that likely accounted for small differences in flagellar angle before and after confinement.

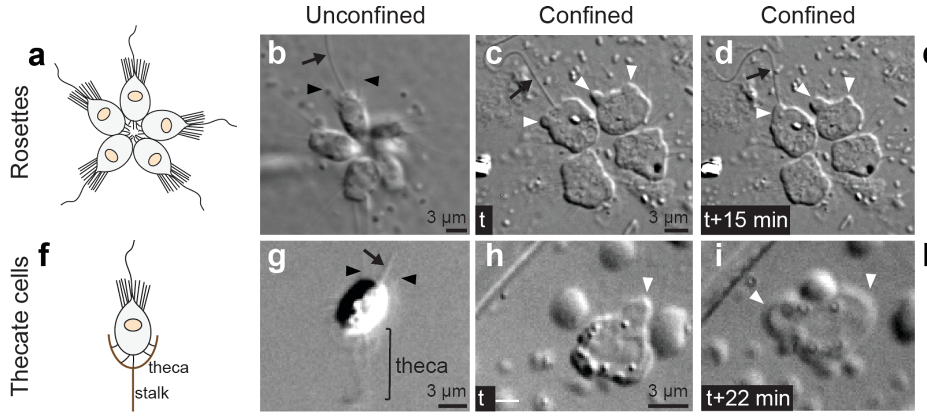


**e**

| Condition | Cells with protrusions | Cells without protrusions |
|---|---|---|
| Unconfined | 0 | 25 |
| Confined | 28 | 0 |

**k**

| Condition | Cells with protrusions | Cells without protrusions |
|---|---|---|
| Unconfined | 0 | 11 |
| Confined | 24 | 0 |

**Extended Data Figure 2. *S. rosetta* is competent to undergo the amoeboid switch in rosette and thecate forms. a,** Schematic drawing of a rosette colony of *S. rosetta* (from[1]). **b-e**, Cells within rosettes became amoeboid under 2 μm confinement. **a**, An unconfined rosette. **c-d**, Time series of a confined rosette, showing dynamic extension and retraction of protrusions. All cells switched to an amoeboid phenotype, but we did not observe any evidence of collective behavior (Fig. 1). **e**, Quantification of the amoeboid switch in rosettes from two biological replicates. Numbers refer to individual cells. **f**, Schematic drawing of a thecate *S. rosetta* cell (from[1]). Thecate cells are sessile and attached to the substrate through an extracellular lodge called a "theca." **g-j**, Thecate cells become amoeboid under 2 μm confinement. **g**, An unconfined thecate cell. **h-i**, A confined thecate cell, showing dynamic extension and retraction of protrusions. **j**, Quantification of the amoeboid switch in a population of confined thecate cells. In all panels, white arrowheads: dynamic protrusions, black arrowheads: microvilli, black arrow: flagellum. Time stamps in black boxes are min:sec.

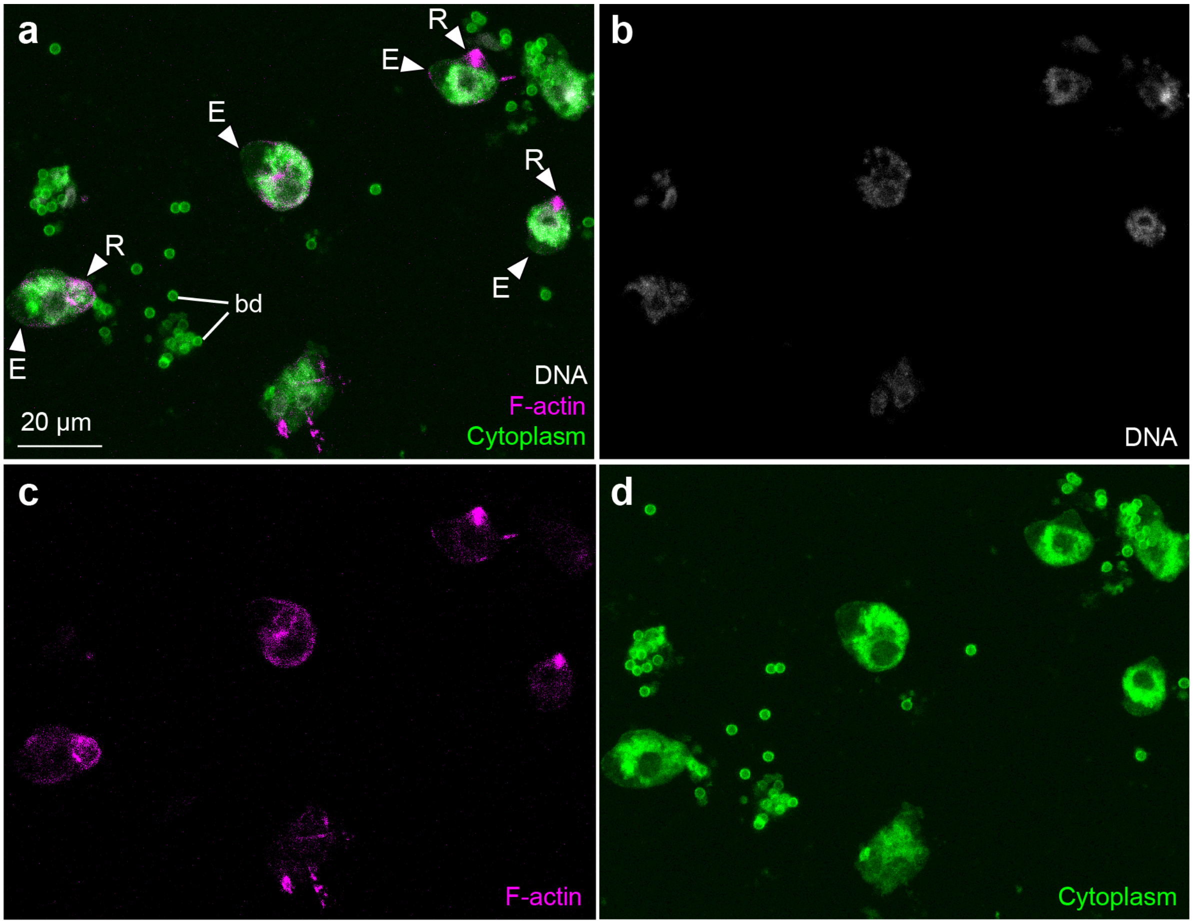


**Extended Data Figure 3. Population of *S. rosetta* cells fixed and stained for F-actin (a, c: phalloidin, magenta), DNA (a, b: Hoechst, white) and cytoplasm (a, d: FM 1-43 FX, which distributes to the cytoplasm of *S. rosetta* following fixation, green).** Note that the cell shown in Fig. 2o-q is present within this field (top right). bd: 1 µm microbeads used as confinement spacers, white arrowheads: blebs, E: expanding blebs, R: retracting blebs.

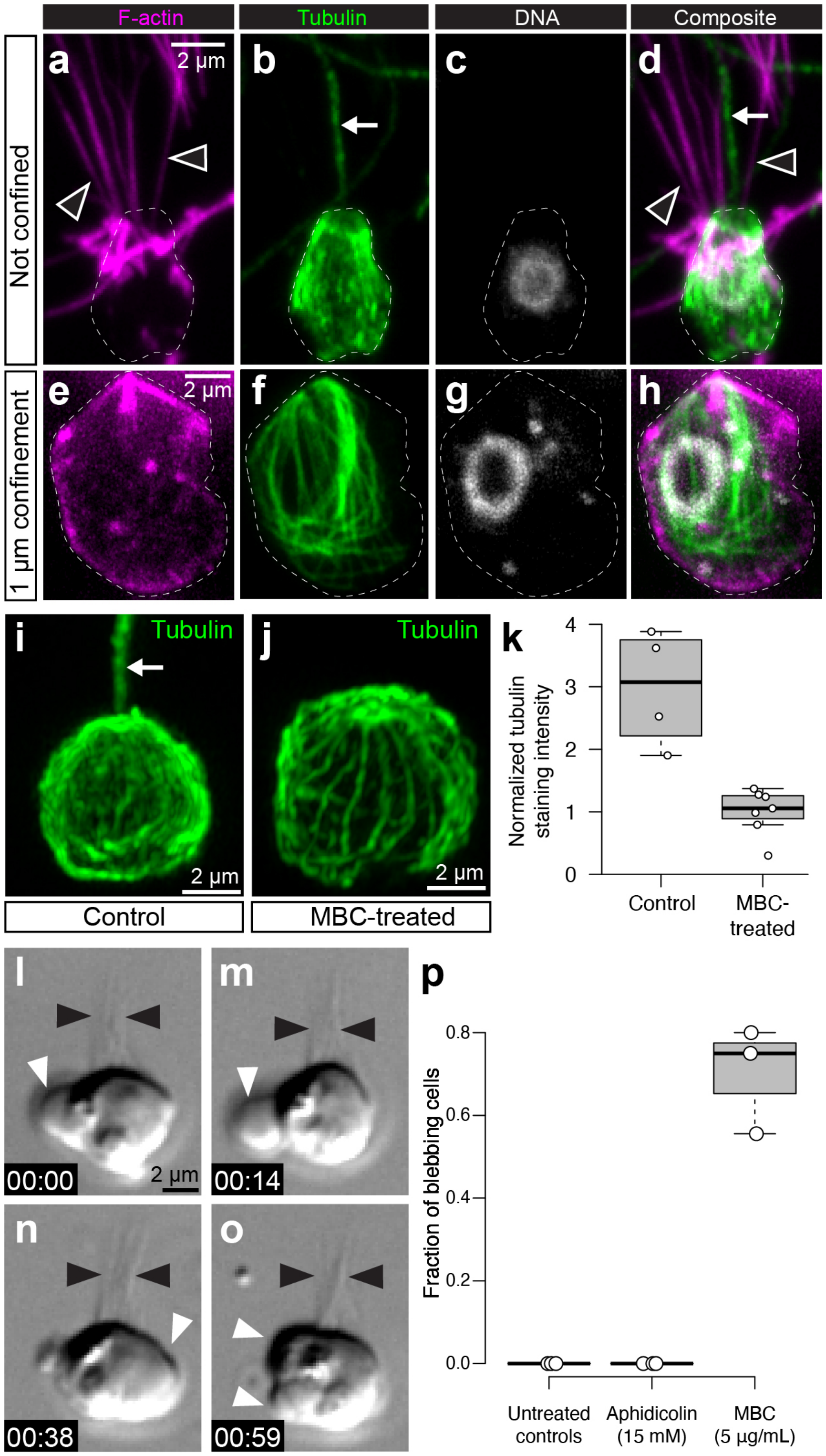
F-actin
Tubulin
DNA
Composite
Not confined
1 µm confinement
a
b
c
d
e
f
g
h
2 µm
i
j
k
Tubulin
Control
MBC-treated
Normalized tubulin staining intensity
0
1
2
3
4
MBC-treated
l
m
n
o
p
00:00
00:14
00:38
00:59
Fraction of blebbing cells
0.0
0.2
0.4
0.6
0.8
Untreated controls
Aphidicolin (15 mM)
MBC (5 µg/mL)

**Extended Data Figure 4. Microtubules regulate blebbing in *S. rosetta*. a-d**, Flagellate *S. rosetta* cells are characterized by a cage of cortical microtubules that underlie the entire plasma membrane, as previously reported[22,62,63]. **a,** A representative flagellated cell, fixed and stained for F-actin (rhodamine-phalloidin, magenta), α-tubulin (YOL3/4 antibody, green) and DNA (Hoechst, white). This immunostained cell was previously published in[40]. **e-h**, Amoeboid *S. rosetta* cells still display a microtubule cage but it is detached from the plasma membrane and surrounds the nucleus. The cell was confined (1 µm), fixed and stained for F-actin (rhodamine-phalloidin, magenta), β-tubulin (E7 antibody, green) and DNA (Hoechst, white). **i-k**, Microtubule depolymerization by treatment with 5 µg/mL MBC (carbendazim) results in loss of the flagellum and reduced cortical microtubule density. **i**, Control cell fixed and stained for β-tubulin (E7 antibody), showing the apical flagellum (arrow) and normal density of cortical microtubules. **j,** MBC-treated cell lacking a flagellum and showing sparser cortical microtubules. **k**, Cortical microtubule density (quantified as normalized β-tubulin staining intensity) is significantly lower in MBC-treated cells than in controls. $p=6.0*10^{-3}$ by the Mann-Whitney U test. **l-o,** Time series of a representative MBC-treated cell forming blebs in the absence of confinement (Supplementary Video 8). **p**, MBC-treated cells reliably form blebs in the absence of confinement, while blebs are never observed in unconfined control cells or in cells treated with aphidicolin (a DNA synthesis inhibitor which does not affect microtubules – but, like MBC, prevents cell division) for the same amount of time. In all panels: white arrowheads: blebs, black arrowheads: microvilli. Time stamps in black boxes are min:sec.

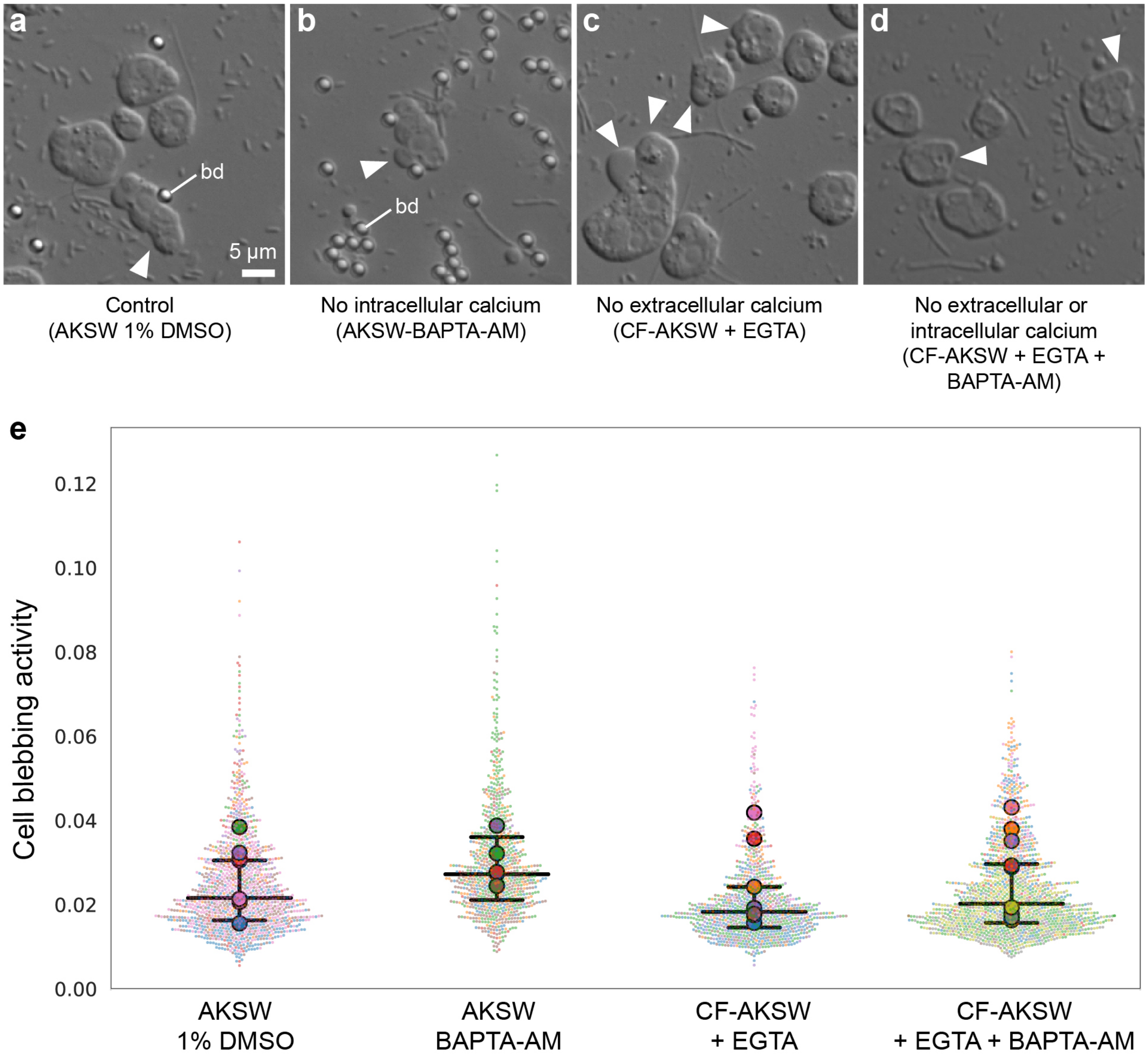


**Extended Data Figure 5. The amoeboid switch is independent of calcium signaling. a-d,** Representative micrographs of *S. rosetta* cells under 2 µm confinement under the following conditions: **a,** in control conditions (AKSW 0.1% DMSO), **b,** after intracellular calcium depletion (with 327 µM of the cell-permeant calcium chelator BAPTA-AM), **c,** after extracellular calcium depletion (calcium-free AKSW (CF-AKSW) with 20 mM of the calcium chelator EGTA), and **d,** after both extracellular and intracellular calcium depletion (CF-AKSW with EGTA and BAPTA-AM). Blebs (white arrowheads) were frequently observed in all conditions. bd: 2 µm microbeads used as spacers. **e**, Quantification of blebbing activity reveals no differences under calcium depletion. Relative blebbing activity of individual cells was quantified by automated cell segmentation and measuring the rate of change of cell shape outline in time (see Methods). Results are depicted as a SuperPlot[98] in which small dots are individual cells and large dots are median values per cell population. Each cell population was treated and imaged as an independent biological replicate. Color of small dots reflects the cell population they belong to. $p$=0.95, 0.91 and 1 respectively for comparisons of the BAPTA-AM, CF-AKSW, and CF-AKSW+BAPTA-AM conditions to the control (by Dunnett's test for comparison of multiple samples to a control). Number of biological replicates and number of cells were respectively: AKSW 1% DMSO control: $N$=7 replicates, $n$=1252 cells; AKSW BAPTA-AM: $N$=6, $n$=921; CF-AKSW: $N$=7, $n$=896; CF-AKSW+BAPTA-AM: $N$=9, $n$=1439.

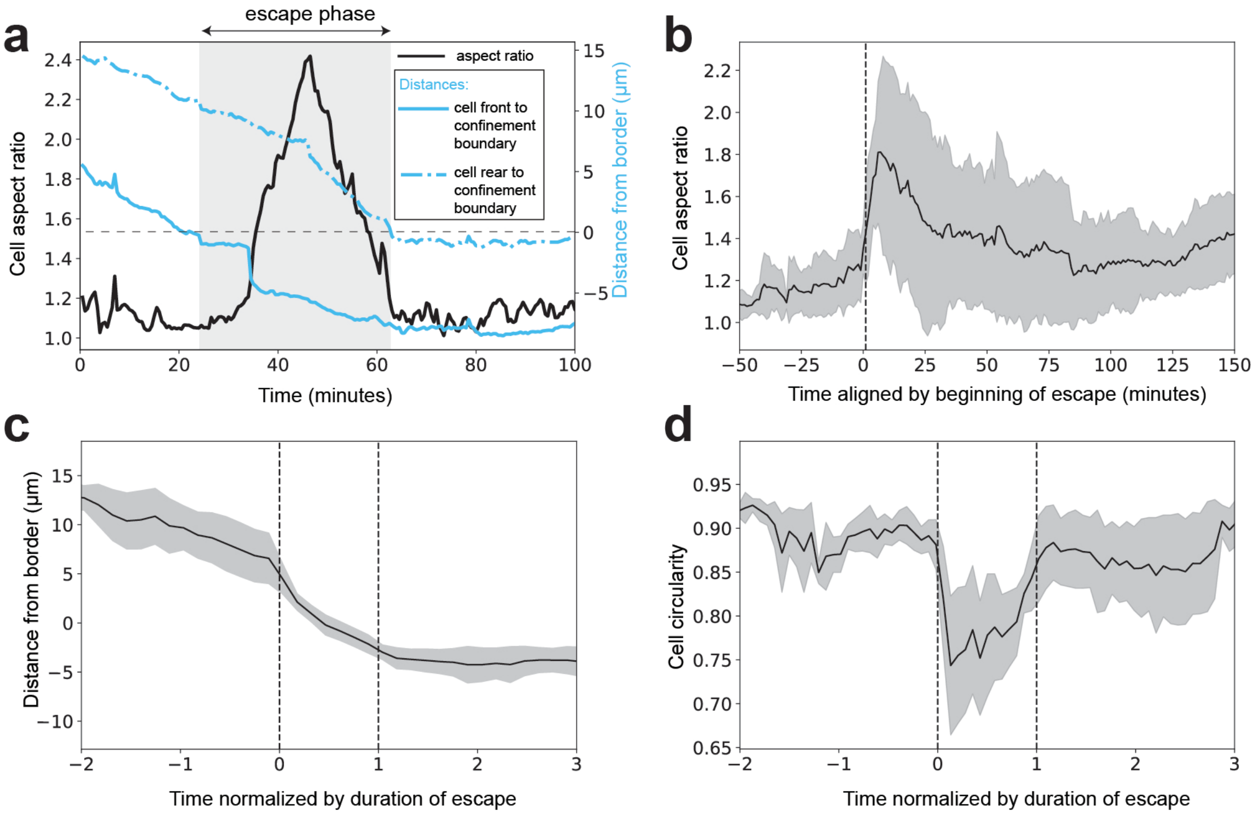


**Extended Data Figure 6. Amoeboid cells elongate during escape from confinement and revert to a rounded shape after escape. a**, Aspect ratio of a representative escaping cell before, during, and after escape (black line). The aspect ratio increased during escape and decreased after escape. Also depicted is the distance between the confinement border and the front or the rear end of the cell (full and dotted blue lines, respectively). The escape phase (in Fig. 3k and panels **c-d** below) is defined and automatically recognized as the time after the front end of the cell crossed the confinement boundary and before the rear end of the cell did. Distance from the border is defined as positive inside the confinement zone and negative in the unconfined space. **b**, Average aspect ratio of escaping cells as a function of absolute time, aligned by beginning of escape. **c**, Distance from the border as a function of time (normalized by duration of the escape phase) in escaping cells. **d,** Cell shape circularity decreased during escape, but resumed a high value once escape was complete, reflecting elongation of the cell during escape (**a**,**b** and Fig. 2k,l). In panels **b-c**, black line is the average value of the metric of interest and grey ribbon is the standard deviation (N=8 escaping cells for all panels).

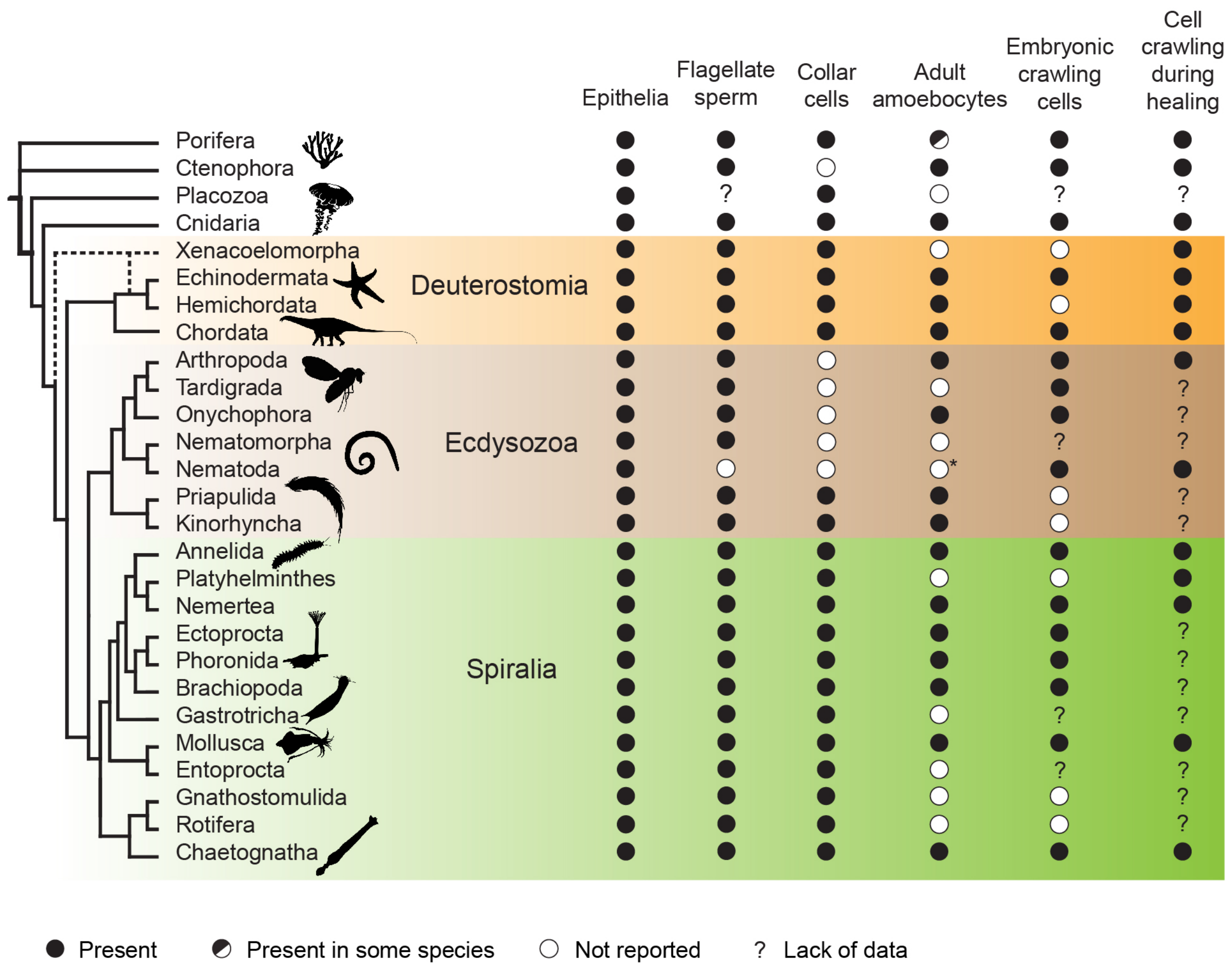


**Extended Data Figure 7. Phylogenetic distribution of crawling cells, epithelial cells, collar cells and flagellated sperm cells in animals.** Shown is a phylogenetic tree (modified from[1]) along with information about the presence or absence (see key) of relevant cell types and cell behaviors in diverse animal lineages. Most animal lineages have crawling cells during both embryonic and adult life history stages. In lineages that lack adult crawling cell types, cell crawling is still often observed in embryonic cells or during wound healing. This phylogenetic distribution suggests that cell crawling was present in the last common animal ancestor – although it might have been restricted to transient developmental or physiological processes (such as primordial germ cell migration or wound healing, respectively), rather than a long-term property of a stable cell type. Cell crawling is seemingly absent from five phyla (Placozoa, Nematomorpha, Gastrotricha, Gnathostomulida and Rotifera), which might reflect lack of data rather than a genuine absence as embryonic development and wound healing are generally little-studied in those groups. Note that amoeboid cell types are thought to be absent in certain lineages within some phyla – such as Calcaronea and possibly Homoscleromorpha within sponges[79]. See Supplementary Table 1 for underlying data and references. Species silhouettes are from Phylopic (http://phylopic.org).

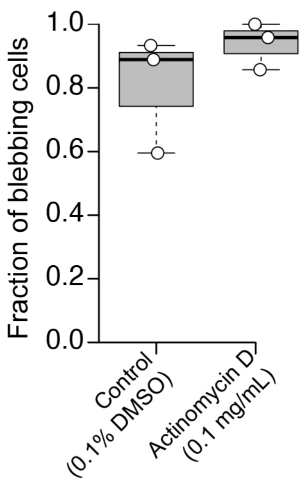


**Extended Data Figure 8. The amoeboid switch is not affected by transcription inhibition.** The fraction of blebbing cells was comparable in DMSO-treated controls (left) and in cells treated with an RNA-polymerase II inhibitor (0.1 mg/mL actinomycin D).

### Supplementary Table 1. Cell crawling across animal diversity.

| Phylum | Crawling cell types in adult organisms | Crawling cells during development | Crawling cells during wound healing |
|---|---|---|---|
| **Porifera** | Archeocytes [1] (absent in calcaroneans [2])<br>Amoeboid sperm in *Leucosolenia* [3] | Transiently amoeboid oocytes [4,5]<br>Ingression during larval development in some species [6] | Epithelial-to-mesenchymal transition [7] or epithelial migration [8] of wounded pinacocytes<br>Crawling of mesohyl cells toward the wound [8] |
| **Ctenophora** | Phagocytic stellate cells [9] | Transiently amoeboid oocytes [10] | Migration of mesogleal cells toward the wound[11] |
| **Placozoa** | *None reported* | *Post-cleavage development unknown*<br>Possible oocyte formation from epithelial cells by epithelial-to-mesenchymal transition and cell migration [12,13] | *Not studied at the cell level* (but see[14] for a tissue-level study of healing) |
| **Cnidaria** | Amoebocytes [15] | Transiently amoeboid oocytes[16–18]<br>Primordial germ cell migration[19]<br>Gastrulation by ingression in some species[20,21] | Epithelial cell migration[22–24]<br>Migration of mesogleal cells toward the wound[25] |
| **Xenacoelomorpha** | *None reported* [26] | *None reported* | Epithelial-to-mesenchymal transition of wounded epidermal cells [27] |
| **Echinodermata** | Coelomocytes [28] | Primordial germ cell migration [29] | Mesenchymal cells migration toward site of injury [30] |
| **Hemichordata** | Coelomocytes [31] | *None reported* (germline development unknown) | Migration toward site of injury [30,32,33] |
| **Chordata** | Amoebocytes in cephalochordates and urochordates [31,34]<br>Leukocytes and mesenchymal cells in vertebrates [35] | Primordial germ cell migration[36–38]<br>Gastrulation by ingression in some species[39]<br>Neural crest cells in vertebrates | Epithelial-to-mesenchymal transition in injured epithelia[40]<br>Leukocyte migration toward site of injury[41–43] |
| **Arthropoda** | Hemocytes [44] | Primordial germ cell migration [45]<br>Gastrulation by ingression in some species [46]<br>Mesenchymal migration of mesodermal cells [47,48] | Epithelial cell migration in injured epithelia [49] |
| **Tardigrada** | *None reported* (so-called brain "amoebocytes" are likely not motile [50]) | Primordial germ cell migration [51,52]<br>Migration of endodermal and mesodermal precursors [51,52] | *Not studied* |
| **Onychophora** | Hemocytes [53] | Primordial germ cell migration [54]<br>Migration of endodermal and mesodermal precursors [54] | *Not studied* |
| **Nematomorpha** | *None reported* | *None reported* (embryonic development is little known) | *Not studied* |

| Phylum | Crawling cell types in adult organisms | Crawling cells during development | Crawling cells during wound healing |
|---|---|---|---|
| **Nematoda** | Amoeboid sperm cells using a non-actin-based mechanism[55] | Actin-based migration of gonad distal tip cells [56]<br>Migration of head mesodermal cells and male linker cell [57] | Epithelial cell migration in embryos[58] but not adults[59] |
| **Priapulida** | Amoebocytes [60,61] | *None reported* | *Not studied* |
| **Kinorhyncha** | Amoebocytes [60,62,63] | *None reported* | *Not studied* |
| **Loricifera** | Coelomocytes [64] | *None reported* | *Not studied* |
| **Annelida** | Coelomocytes [65] and hemocytes [44] | Primordial germ cell migration [66] | Migration during regeneration [67] |
| **Platyhelminthes** | *None reported* | *None reported* | Neoblasts migrate to the wounded area upon injury [68] |
| **Nemertea** | Hemocytes [69] | Transiently amoeboid oocytes [70]<br>Migratory proboscis precursors [71] | Migration during regeneration [72] |
| **Ectoprocta** | Amoebocytes [73] | Primordial germ cell migration [74] | *Not studied* |
| **Phoronida** | Amoebocytes [75,76] | Migration of mesodermal cells [77,78] | *Not studied* |
| **Brachiopoda** | Amoebocytes [79,80] | *None reported* | *Not studied* |
| **Mollusca** | Hemocytes [44,81] | Transiently amoeboid oocytes [82–85]<br>Presumptive germ cell migration [86]<br>Ectomesoderm migration [87] | Hemocyte migration toward site of injury[88] |
| **Entoprocta** | *None reported* | *None reported* | *Not studied* |
| **Gastrotricha** | *None reported* | *None reported* | *Not studied* |
| **Gnathostomulida** | *None reported* | *None reported* | *Not studied* |
| **Rotifera** | *None reported* | *None reported* | *Not studied* |
| **Micrognathozoa** | *None reported* | *None reported* | *Not studied* |
| **Chaetognatha** | Amoebocytes [89] | Primordial germ cells migration [90] | Epithelial cell migration [91] |

**References associated to Supplementary Table 1**


1. Brusca, R. C. & Brusca, G. J. *Invertebrates*. (Sinauer Associates, Inc., 2003).
2. Adamska, M. Sponges as models to study emergence of complex animals. *Current Opinion in Genetics and Development* (2016). doi:10.1016/j.gde.2016.05.026
3. Anakina, R. P. & Drozdov, A. L. Gamete Structure and Fertilization in the Barents Sea Sponge *Leucosolenia complicata*. *Russ. J. Mar. Biol.* (2001). doi:10.1023/A:1016761317637
4. Franzen, W. Oogenesis and larval development of *Scypha ciliata* (Porifera, Calcarea). *Zoomorphology* **107**, 349–357 (1988).
5. Haeckel, E. *The evolution of man: a popular exposition of the principal points of human ontogeny and phylogeny*. (1874). doi:10.5962/bhl.title.61275
6. Ereskovsky, A. V. *The comparative embryology of sponges*. *The Comparative Embryology of Sponges* (Springer, 2010). doi:10.1007/978-90-481-8575-7
7. Borisenko, I. E., Adamska, M., Tokina, D. B. & Ereskovsky, A. V. Transdifferentiation is a driving force of regeneration in *Halisarca dujardini* (Demospongiae, Porifera). *PeerJ* (2015). doi:10.7717/peerj.1211
8. Lavrov, A. I., Bolshakov, F. V., Tokina, D. B. & Ereskovsky, A. V. Sewing up the wounds : The epithelial morphogenesis as a central mechanism of calcaronean sponge regeneration. *J. Exp. Zool. Part B Mol. Dev. Evol.* (2018). doi:10.1002/jez.b.22830
9. Traylor-Knowles, N., Vandepas, L. E. & Browne, W. E. Still Enigmatic: Innate Immunity in the Ctenophore *Mnemiopsis leidyi*. in *Integrative and Comparative Biology* (2019). doi:10.1093/icb/icz116
10. Dunlap-Pianka, H. Ctenophora. in *Reproduction of Marine Invertebrates, volume I. Acoelomate and Pseudocoelomate Metazoans* (eds. Giese, A. C. & Pearse, J. S.) (Academic Press, 1974). doi:10.1016/b978-0-12-282501-9.50009-0
11. Ramon-Mateu, J., Ellison, S. T., Angelini, T. E. & Martindale, M. Q. Regeneration in the ctenophore

*Mnemiopsis leidyi* occurs in the absence of a blastema, requires cell division, and is temporally separable from wound healing. *BMC Biol.* (2019). doi:10.1186/s12915-019-0695-8

12. Grell, K. G. & Ruthmann, A. Placozoa. in *Microscopic Anatomy of Invertebrates. Volume 2: Placozoa, Porifera, Cnidaria, and Ctenophora* 13–28 (Wiley-Liss, Inc., 1991).
13. Grell, K. G. & Benwitz, G. Elektronenmikroskopische beobachtungen über das wachstum der eizelle und die bildung der 'befruchtungsmembran' von *Trichoplax adhaerens* F. E. Schulze (Placozoa). *Zeitschrift für Morphol. der Tiere* (1974). doi:10.1007/BF00277511
14. Prakash, V. N., Bull, M. S. & Prakash, M. Motility induced fracture reveals a ductile to brittle crossover in the epithelial tissues of a simple animal. *bioRxiv* (2019). doi:10.1101/676866
15. Gold, D. A. & Jacobs, D. K. Stem cell dynamics in Cnidaria: Are there unifying principles? *Dev. Genes Evol.* (2013). doi:10.1007/s00427-012-0429-1
16. Honegger, T. G., Zürrer, D. & Tardent, P. Oogenesis in *Hydra carnea*: A new model based on light and electron microscopic analyses of oocyte and nurse cell differentiation. *Tissue Cell* (1989). doi:10.1016/0040-8166(89)90052-9
17. Larkman, A. U. An ultrastructural study of oocyte growth within the endoderm and entry into the mesoglea in *Actinia fragacea* (Cnidaria, anthozoa). *J. Morphol.* (1983). doi:10.1002/jmor.1051780207
18. Eckelbarger, K. J., Hand, C. & Uhlinger, K. R. Ultrastructural features of the trophonema and oogenesis in the starlet sea anemone, *Nematostella vectensis* (Edwardsiidae). *Invertebr. Biol.* (2008). doi:10.1111/j.1744-7410.2008.00146.x
19. Chen, C.-Y., McKinney, S. A., Ellington, L. R. & Gibson, M. C. Hedgehog signaling is required for endomesodermal patterning and germ cell development in *Nematostella vectensis*. *bioRxiv* (2020). doi:10.1101/2020.01.15.907238
20. Kraus, Y., Chevalier, S. & Houliston, E. Cell shape changes during larval body plan development in *Clytia hemisphaerica*. *bioRxiv* (2019). doi:10.1101/864223
21. Martindale, M. Q. & Byrum, C. A. Gastrulation in the Cnidaria and Ctenophora. in *Gastrulation* (ed. Stern, C. D.) 731 (Cold Spring Harbor Laboratory Press, 2004).
22. Malamy, J. E. & Shribak, M. An orientation-independent DIC microscope allows high resolution imaging of epithelial cell migration and wound healing in a cnidarian model. *J. Microsc.* (2018). doi:10.1111/jmi.12682
23. Kamran, Z. *et al.* In vivo imaging of epithelial wound healing in the cnidarian *Clytia hemisphaerica* demonstrates early evolution of purse string and cell crawling closure mechanisms. *BMC Dev. Biol.* (2017). doi:10.1186/s12861-017-0160-2
24. DuBuc, T. Q., Traylor-Knowles, N. & Martindale, M. Q. Initiating a regenerative response; cellular and molecular features of wound healing in the cnidarian *Nematostella vectensis*. *BMC Biol.* (2014). doi:10.1186/1741-7007-12-24
25. Patterson, M. J. & Landolt, M. L. Cellular reaction to injury in the anthozoan *Anthopleura elegantissima*. *J. Invertebr. Pathol.* (1979). doi:10.1016/0022-2011(79)90152-6
26. Chiodin, M. *et al.* Mesodermal Gene Expression in the Acoel Isodiametra pulchra Indicates a Low Number of Mesodermal Cell Types and the Endomesodermal Origin of the Gonads. *PLoS One* (2013). doi:10.1371/journal.pone.0055499
27. Geddes, P. Observations on the Physiology and Histology of Convoluta Schultzii. *Proc. R. Soc. London* **28**, 449–457 (1879).
28. Chia, F. S. & Xing, J. Echinoderm coelomocytes. *Zoological Studies* (1996).
29. Campanale, J. P. *et al.* Migration of sea urchin primordial germ cells. *Dev. Dyn.* (2014). doi:10.1002/dvdy.24133
30. Rychel, A. L. & Swalla, B. J. Regeneration in hemichordates and echinoderms. in *Stem Cells in Marine Organisms* (2009). doi:10.1007/978-90-481-2767-2_10
31. Rowley, A. F., Rhodes, C. P. & Ratcliffe, N. A. Protochordate leucocytes: a review. *Zool. J. Linn. Soc.* (1984). doi:10.1111/j.1096-3642.1984.tb01978.x
32. Rychel, A. L. & Swalla, B. J. Anterior regeneration in the hemichordate Ptychodera flava. *Dev. Dyn.* (2008). doi:10.1002/dvdy.21747
33. Rao, K. P. Morphogenesis during regeneration is an enteropneust. *J. Anim. Morphol. Physiol. India* 1–7 (1955).
34. Muñoz-Chápuli, R. & Pérez-Pomares, J. A. M. Origin of the Vertebrate Endothelial Cell Lineage. Ontogeny and Phylogeny. in *Heart Development and Regeneration* (2010). doi:10.1016/B978-0-12-381332-9.00022-0
35. Kierszenbaum, A. L. & Tres, L. L. Blood and hematopoiesis. in *Histology and Cell Biology: An Introduction to Pathology* 752 (Saunders, 2015). doi:10.1016/b978-0-323-07842-9.50010-1

36. Grimaldi, C. & Raz, E. Germ cell migration—Evolutionary issues and current understanding. *Seminars in Cell and Developmental Biology* (2020). doi:10.1016/j.semcdb.2019.11.015
37. Tarbashevich, K. & Raz, E. The nuts and bolts of germ-cell migration. *Current Opinion in Cell Biology* (2010). doi:10.1016/j.ceb.2010.09.005
38. Nieuwkoop, P. D. & Sutasurya, L. A. *Primordial germ cells in the chordates : embryogenesis and phylogenesis*. *Developmental and cell biology series* (Cambridge University Press, 1979).
39. Shook, D. R. & Keller, R. Epithelial type, ingression, blastopore architecture and the evolution of chordate mesoderm morphogenesis. *Journal of Experimental Zoology Part B: Molecular and Developmental Evolution* (2008). doi:10.1002/jez.b.21198
40. Stone, R. C. *et al.* Epithelial-mesenchymal transition in tissue repair and fibrosis. *Cell and Tissue Research* (2016). doi:10.1007/s00441-016-2464-0
41. Powell, D. *et al.* Chemokine Signaling and the Regulation of Bidirectional Leukocyte Migration in Interstitial Tissues. *Cell Rep.* (2017). doi:10.1016/j.celrep.2017.04.078
42. De Oliveira, S., Rosowski, E. E. & Huttenlocher, A. Neutrophil migration in infection and wound repair: Going forward in reverse. *Nature Reviews Immunology* (2016). doi:10.1038/nri.2016.49
43. Sieger, D., Moritz, C., Ziegenhals, T., Prykhozhij, S. & Peri, F. Long-Range Ca2+ Waves Transmit Brain-Damage Signals to Microglia. *Dev. Cell* (2012). doi:10.1016/j.devcel.2012.04.012
44. Stang-Voss, C. On the Ultrastructure of Invertebrate Hemocytes: An Interpretation of their Role in Comparative Hematology. in *Contemporary Topics in Immunobiology, volume 4: Invertebrate Immunology* (ed. Cooper, E. L.) 65–76 (Plenum Press, New York, 1974). doi:10.1007/978-1-4684-3048-6_7
45. Warrior, R. Primordial germ cell migration and the assembly of the *Drosophila* embryonic gonad. *Dev. Biol.* (1994). doi:10.1006/dbio.1994.1306
46. Urbansky, S., Avalos, P. G., Wosch, M. & Lemke, S. Folded gastrulation and T48 drive the evolution of coordinated mesoderm internalization in flies. *Elife* (2016). doi:10.7554/eLife.18318.001
47. Smallhorn, M., Murray, M. J. & Saint, R. The epithelial-mesenchymal transition of the *Drosophila* mesoderm requires the Rho GTP exchange factor pebble. *Development* (2004). doi:10.1242/dev.01150
48. Schäfer, G., Narasimha, M., Vogelsang, E. & Leptin, M. Cadherin switching during the formation and differentiation of the *Drosophila* mesoderm - implications for epithelial-tomesenchymal transitions. *J. Cell Sci.* (2014). doi:10.1242/jcs.139485
49. Wu, Y. *et al.* A Blood-Borne PDGF/VEGF-like Ligand Initiates Wound-Induced Epidermal Cell Migration in *Drosophila* Larvae. *Curr. Biol.* (2009). doi:10.1016/j.cub.2009.07.019
50. Persson, D. K., Halberg, K. A., Jørgensen, A., Møbjerg, N. & Kristensen, R. M. Brain anatomy of the marine tardigrade *Actinarctus doryphorus* (Arthrotardigrada). *J. Morphol.* (2014). doi:10.1002/jmor.20207
51. Hejnol, A. & Schnabel, R. The eutardigrade *Thulinia stephaniae* has an indeterminate development and the potential to regulate early blastomere ablations. *Development* (2005). doi:10.1242/dev.01701
52. Hejnol, A. & Schnabel, R. What a couple of dimensions can do for you: Comparative developmental studies using 4D microscopy - Examples from tardigrade development. *Integr. Comp. Biol.* (2006). doi:10.1093/icb/icj012
53. Silva, J. R. M. C., Coelho, M. P. D. & Nogueira, M. I. Induced inflammatory process in Peripatus acacioi Marcus et Marcus (Onychophora). *J. Invertebr. Pathol.* (2000). doi:10.1006/jipa.1999.4898
54. Anderson, D. T. *Embryology and Phylogeny in Annelids and Arthropods.* (Pergamon Press, 1973). doi:10.2307/2412250
55. Roberts, T. M. & Stewart, M. Nematode sperm: Amoeboid movement without actin. *Trends in Cell Biology* (1997). doi:10.1016/S0962-8924(97)01113-6
56. Cram, E. J., Shang, H. & Schwarzbauer, J. E. A systematic RNA interference screen reveals a cell migration gene network in *C. elegans*. *J. Cell Sci.* (2006). doi:10.1242/jcs.03274
57. Hedgecock, E. M., Culotti, J. G. & Hall, D. H. The unc-5, unc-6, and unc-40 genes guide circumferential migrations of pioneer axons and mesodermal cells on the epidermis in *C. elegans*. *Neuron* (1990). doi:10.1016/0896-6273(90)90444-K
58. Raich, W. B., Agbunag, C. & Hardin, J. Rapid epithelial-sheet sealing in the *Caenorhabditis elegans* embryo requires cadherin-dependent filopodial priming. *Curr. Biol.* (1999). doi:10.1016/S0960-9822(00)80015-9
59. Xu, S. & Chisholm, A. D. A Gαq-Ca2+ signaling pathway promotes actin-mediated epidermal wound closure in *C. elegans*. *Curr. Biol.* (2011). doi:10.1016/j.cub.2011.10.050
60. Schmidt-Rhaesa, A. *The Evolution of Organ Systems*. *The Evolution of Organ Systems* (2007). doi:10.1093/acprof:oso/9780198566687.001.0001

61. McLean, N. Amoebocytes in the Lining of the Body Cavity and Mesenteries of *Priapulus caudatus* (Priapulida). *Acta Zool.* (1984). doi:10.1111/j.1463-6395.1984.tb00811.x
62. Kristensen, R. M. & Hay-Schmidt, A. The Protonephridia of the Arctic Kinorhynch *Echinoderes aquilonius* (Cyclorhagida, Echinoderidae). *Acta Zool.* (1989). doi:10.1111/j.1463-6395.1989.tb01048.x
63. Neuhaus, B. & Higgins, R. P. Ultrastructure, biology, and phylogenetic relationships of Kinorhyncha. in *Integrative and Comparative Biology* (2002). doi:10.1093/icb/42.3.619
64. Kristensen, R. M. Loricifera. in *Microscopic Anatomy of Invertebrates. Volume 4: Aschelminthes* 448 (Wiley-Liss, Inc., 1991).
65. Baskin, D. G. The Coelomocytes of Nereid Polychaetes. in *Contemporary Topics in Immunobiology, volume 4: Invertebrate Immunology* (ed. Cooper, E. L.) 55–64 (Plenum Press, New York, 1974). doi:10.1007/978-1-4684-3048-6_6
66. Rebscher, N., Zelada-González, F., Banisch, T. U., Raible, F. & Arendt, D. Vasa unveils a common origin of germ cells and of somatic stem cells from the posterior growth zone in the polychaete *Platynereis dumerilii*. *Dev. Biol.* (2007). doi:10.1016/j.ydbio.2007.03.521
67. Zattara, E. E., Turlington, K. W. & Bely, A. E. Long-term time-lapse live imaging reveals extensive cell migration during annelid regeneration. *BMC Dev. Biol.* (2016). doi:10.1186/s12861-016-0104-2
68. Palmberg, I. Cell migration and differentiation during wound healing and regeneration in *Microstomum lineare* (Turbellaria). in *Advances in the Biology of Turbellarians and Related Platyhelminthes* (1986). doi:10.1007/978-94-009-4810-5_25
69. Valembois, P., Roch, P. & Boiledieu, D. Cellular Defense Systems of the Platyhelminthes, Nemertea, Sipunculida, and Annelida. in *The Reticuloendothelial System: A Comprehensive Treatise. Volume 3: Phylogeny and Ontogeny* 773 (Plenum Press, New York, 1982). doi:10.1007/978-1-4684-4166-6_4
70. Crandall, F. B., Norenburg, J. L. & Gibson, R. Gonadogenesis, embryogenesis, and unusual oocyte origin in *Notogaeanemertes folzae* Riser, 1988 (Nemertea, Hoplonemertea). in *Hydrobiologia* (1997). doi:10.1023/A:1003145519006
71. Smith, J. E. Memoirs: The Early Development of the Nemertean *Cephalothrix rufifrons*. *J. Cell Sci.* **2**, 335–381 (1935).
72. Coe, W. R. Analysis of the regenerative processes in nemerteans. *Biol. Bull.* **66**, 304–315 (1934).
73. Xing, J. & Qian, P. Y. Tower cells of the marine bryozoan *Membranipora membranacea*. *J. Morphol.* (1999). doi:10.1002/(SICI)1097-4687(199902)239:2<121::AID-JMOR1>3.0.CO;2-1
74. Dyrynda, P. E. J. & King, P. E. Gametogenesis in placental and non-placental ovicellate cheilostome Bryozoa. *J. Zool.* (1983). doi:10.1111/j.1469-7998.1983.tb02810.x
75. Temereva, E. N. & Malakhov, V. V. The circulatory system of phoronid larvae. *Dokl. Biol. Sci.* **375**, 712–714 (2000).
76. Pardos, F., Roldán, C., Benito, J. & Emig, C. C. Fine Structure of the Tentacles of *Phoronis australis* Haswell (Phoronida, Lophophorata). *Acta Zool.* (1991). doi:10.1111/j.1463-6395.1991.tb00320.x
77. Malakhov, V. V. & Temereva, E. N. Embryonic Development of the Phoronid *Phoronis ijimai*. *Russ. J. Mar. Biol.* (2000). doi:10.1023/A:1009494621160
78. Temereva, E. n. & Malakhov, V. v. Embryogenesis and larval development of *Phoronopsis harmeri* Pixell, 1912 (Phoronida): Dual origin of the coelomic mesoderm. *Invertebr. Reprod. Dev.* (2007). doi:10.1080/07924259.2007.9652228
79. Rowley, A. F. & Hayward, P. J. Blood cells and coelomocytes of the inarticulate brachiopod *Lingula anatina*. *J. Zool.* (1985). doi:10.1111/j.1469-7998.1985.tb05609.x
80. Kuzmina, T. V., Temereva, E. N. & Malakhov, V. V. Ultrastructure of the lophophoral coelomic lining in the brachiopod *Hemithiris psittacea*: functional and evolutionary significance. *Zoomorphology* (2018). doi:10.1007/s00435-018-0397-8
81. Sminia, T. Phagocytic Cells in Molluscs. in *Aspects of Developmental and Comparative Immunology* (1981). doi:10.1016/b978-0-08-025922-2.50022-x
82. Saleuddin, A. S. M. & Khan, H. R. Motility of the oocyte of *Helisoma* (Mollusca). *Eur. J. Cell Biol.* (1981).
83. Saleuddin, A. S. M. & Farrell, C. L. Brain extract causes amoeboid movement in vitro in oocytes in Helix aspersa (Mollusca). *Int. J. Invertebr. Reprod.* (1983). doi:10.1080/01651269.1983.10510021
84. Bretschneider, L. H. & Raven, C. P. Structural and Topochemical Changes in the Egg Cells of *Limnaea stagnalis* L. During Oogenesis. *Netherlands J. Zool.* (1954). doi:10.1163/036551654X00113
85. Blaauw-Jansen, G. On the influence of temperature and of lithium chloride on the amoeboid mobility of unsegmented eggs of *Limnaea stagnalis* L. *Proc. K. Ned. Akad. van Wet.* **53**, 910–912 (1950).
86. Lyons, D. C., Perry, K. J., Lesoway, M. P. & Henry, J. Q. Cleavage pattern and fate map of the

mesentoblast, 4d, in the gastropod *Crepidula*: a hallmark of spiralian development. *Evodevo* 21 (2012).
87. Lyons, D. C., Perry, K. J. & Henry, J. Q. Spiralian gastrulation: Germ layer formation, morphogenesis, and fate of the blastopore in the slipper snail *Crepidula fornicata*. *Evodevo* (2015). doi:10.1186/s13227-015-0019-1
88. Franchini, A. & Ottaviani, E. Repair of molluscan tissue injury: Role of PDGF and TGF-β. *Tissue Cell* (2000). doi:10.1054/tice.2000.0118
89. Malakhov, V. V. & Berezinskaya, T. L. Structure of the Circulatory System of Arrow Worms (Chaetognatha). *Dokl. Biol. Sci.* **376**, 78–80 (2001).
90. Carré, D., Djedlat, C. & Sardet, C. Formation of a large Vasa-positive germ granule and its inheritance by germ cells in the enigmatic Chaetognaths. *Development* (2002).
91. Duvert, M., Perez, Y. & Casanova, J. P. Wound healing and survival of beheaded chaetognaths. *J. Mar. Biol. Assoc. United Kingdom* (2000). doi:10.1017/S0025315400002873

### Supplementary Videos

**Video 1.** Time-lapse of an *S. rosetta* cell undergoing progressive confinement by evaporation and switching to an amoeboid phenotype. The strain used was SrEpac and the starting cell type was slow swimmer.

**Video 2.** Time-lapse of a population of *S. rosetta* cells before and during 2 µm confinement under a confinement slide controlled by a dynamic cell confiner. The strain used was SrEpac and the starting cell type was slow swimmer.

**Video 3.** Time-lapse of a population of *S. rosetta* cells before, during and after 2 µm confinement under a confinement slide controlled by a dynamic cell confiner. Cells were attached to the substrate by poly-D-lysine to minimize cell movement and help visualizing both conversion of flagellates into amoeboid cells, and reversion of amoeboid cells back into flagellates. For this reason, cells show less crawling movements (compare Supplementary Video 2). The strain used was SrEpac and the starting cell type was slow swimmer.

**Video 4.** Time-lapse of a population of *S. rosetta* cells confined between two glass cover slips using 2 µm microbeads as spacers. The strain used was SrEpac and the starting cell type was slow swimmer.

**Video 5.** Time-lapse of a population of *S. rosetta* cells confined in a thinly spread liquid film under a layer of anti-evaporation oil (see Methods). The strain used was SrEpac and the starting cell type was slow swimmer.

**Video 6.** Time-lapse of a population of *S. rosetta* cells confined on the surface of a 1% agar gel in artificial seawater, under a layer of anti-evaporation oil. The strain used was SrEpac and the starting cell type was slow swimmer.

**Video 7.** Time-lapse of an *S. rosetta* cell transfected with LifeAct-mCherry (which marks F-actin) and septin2-mTFP (which here distributed into most of the cytoplasm). Blebs first form as cytoplasm-filled, F-actin-free protrusions, and are re-invaded by F-actin before retraction.

**Video 8.** Time-lapse of an *S. rosetta* cell treated with the microtubule-depolymerizing drug MBC.

**Video 9.** Time-lapse of a crawling *S. rosetta* cell transfected with LifeAct-mCherry and septin2-mTFP. Note F-actin retrograde flow within the leading bleb (Fig. 3g-h).

**Video 10.** Time-lapse of an mTFP-expressing population of *S. rosetta* cells trapped in a 0.5 μm space under a circular micropillar. Some of the most peripheral cells (<10 μm distant of the border) manage to cross the border into the unconfined space around the pillar.

**Video 11.** Close-up of an escaping cell (from Supplementary Video 10) showing DIC channel (top left), mTFP channel (top right), segmented cell shape (bottom left) and segmented cell shape (magenta) overlaid with the DIC channel (grey). The cell elongates and polarizes in the direction of the border during crossing, and resumes a round shape after having crossed.

**Video 12.** Time-lapse of an escaping cell (from a similar experiment to the one shown in Supplementary Video 10) shedding two blebs into the unconfined space prior to escaping. One of the blebs is then reabsorbed by the cell during escape from confinement.

**Video 13.** Time-lapse of a 2 μm-confined choanoflagellate of the species *Diaphanoeca grandis*. The cage surrounding the cell is called a "lorica" and is a basket of silicon strips secreted and assembled by the cell. The cell is flattened but does not show blebbing or active deformation.

**Video 14.** Time-lapse of a population of *Monosiga brevicollis* confined in a thin liquid film, displaying intense blebbing and crawling (or gliding) motility.

**Video 15.** Time-lapse of a 2 µm-confined *Acanthoeca spectabilis* showing dynamic bleb extension.

**Video 16.** Time-lapse of four *Salpingoeca helianthica* cells, 2 µm-confined, displaying long dynamic blebs.

**Video 17.** Time-lapse of a 2 µm-confined *Salpingoeca urceolata* cell crawling out of its theca. The cell crawls over about 40 µm, shedding cellular material at its rear end (possibly similar to the shedding of blebs by *S. rosetta*; Supplementary Video 12).**q**

**Video 18.** Automated recognition of blebs in 2 µm-confined *S. rosetta* cells. Left panel: DIC channel. Middle panel: result of the cell segmentation superimposed onto the DIC channel. Right panel: cell protrusions, classified into expanding blebs (orange) and retracting blebs (blue).
